# δ2-Protocadherins organize parallel indirect basal ganglia circuits

**DOI:** 10.64898/2025.12.13.694147

**Authors:** Naosuke Hoshina, Joshua M. Boeckers, Erin M. Johnson-Venkatesh, Miyuki Hoshina, Kana Matsumoto, Abhijnana Das, Veronica R. Rally, Jaanvi Sant, Takafumi Inoue, Hisashi Umemori

## Abstract

The basal ganglia (BG) contain multiple parallel neural circuits, each of which may control different behaviors. However, how the distinct parallel BG circuits are molecularly organized is not known. Here we show that two δ2-protocadherins (PCDHs), PCDH17 and PCDH10, which are homophilic cell-adhesion molecules, establish and define two distinct indirect BG circuits that regulate different behaviors. PCDH17 and PCDH10 are expressed in a complementary expression pattern in the BG, anatomically defining two parallel indirect BG connections. Indirect pathway-specific *Pcdh17* and *Pcdh10* conditional knockout (cKO) mice show impaired establishment of the indirect BG circuits in a region-preferential manner. Finally, the *Pcdh17*-cKO mice show defects in task learning, while the *Pcdh10*-cKO mice show defects in motor/sensory habituation. These results identify PCDH17 and PCDH10 as the molecular organizers for two distinct indirect BG circuits regulating different behaviors and reveal the molecular mechanisms for organizing parallel BG circuits.

**Teaser:** Distinct protocadherins organize parallel indirect basal ganglia circuits that regulate task learning or sensorimotor habituation

## Introduction

The basal ganglia (BG) comprise a group of subcortical nuclei that include the striatum, external globus pallidus (GPe), internal globus pallidus (GPi), substantia nigra pars reticulata (SNr), and subthalamic nucleus (STN). The striatum receives input from the cerebral cortex and projects to the GPi/SNr directly (direct pathway) or indirectly through the GPe and STN (indirect pathway). The GPi/SNr sends output back to the cortex via the thalamus, thereby constituting the cortico-BG-thalamo-cortical loop. Different cortical areas project to discrete regions of the BG in a topographic manner, constituting a parallel organization of functionally segregated circuits (*1–3*). Neurons in the ventromedial prefrontal and orbitofrontal cortices project to the ventral striatum; the dorsolateral prefrontal cortex to the anterior striatum; and the motor cortex to more caudal regions of the striatum, forming limbic, associative, and sensorimotor circuits, respectively (*4*, *5*). The limbic circuit appears to play a role in motivated behavior and empathic/socially appropriate behavior; the associative circuit is implicated in executive and cognitive function; and the sensorimotor circuit plays a role in sensorimotor modulation and movement regulation. While the anatomical distribution of parallel cortico-BG circuits has been described in mice (*6*), we do not know how these distinct circuits are organized molecularly - we do not know the molecular mechanisms by which these parallel BG circuits are established. Defects in the cortico-BG circuits are linked to many neuropsychiatric disorders such as Parkinson’s disease, autism spectrum disorder, obsessive-compulsive disorder, schizophrenia, and depression (*1*, *7*). Thus, understanding how parallel cortico-BG circuits are organized should provide insight into potential strategies for treating the particular symptoms of each neuropsychiatric disorder.

Cell adhesion molecules play important roles in connecting appropriate neurons in the brain. For example, anatomically segregated parallel hippocampal circuits are organized based on the expression of distinct cell adhesion molecules (*8*). We have recently found that two members of the δ2-Protocadherin (PCDH) family (*9*), PCDH17 and PCDH10, are expressed in a complementary pattern in the BG (*10*), with PCDH17 and PCDH10 expression in regions implicated in associative and limbic circuits, respectively. δ2-PCDHs are homophilic cell adhesion molecules, suggesting that they may serve as molecular organizers for the establishment of parallel BG circuits.

In this study, we focused on the striatal indirect pathway and examined the roles of PCDH17 and PCDH10 in the establishment of parallel BG circuits. PCDH17- and PCDH10-positive indirect spiny projection neurons (iSPNs) show complementary axonal projection patterns in the GPe that reflect parallel BG circuits. Indirect pathway-specific *Pcdh17* and *Pcdh10* conditional knockout (cKO) mice exhibit aberrant cell patterning and synapse development in the BG in a region-preferential manner. Behaviorally, the *Pcdh17* cKO mice show deficits in task learning, while the *Pcdh10* cKO mice show defects in locomotor and sensory habituation. These results show that parallel BG circuits are molecularly defined and that PCDH17 and PCDH10 organize functionally segregated, parallel indirect BG circuits.

## Results

### PCDH17 and PCDH10 define parallel iSPN axonal connections

PCDH17 and PCDH10 are closely related PCDH members, which are transmembrane proteins with six extracellular cadherin domains. Both proteins mediate highly selective homophilic interactions, but not heterophilic interactions with each other (*10*). We first examined their expression patterns in the BG. Double staining of BG sections for PCDH17 and PCDH10 proteins showed complementary expression patterns in the BG subregions, including the striatum, GPe, GPi, and SNr (Fig. 1A, fig. S1 and fig. S2). PCDH17 protein is mainly expressed in the anterior striatum, whereas PCDH10 protein is preferentially expressed in the posterior striatum, showing a complementary expression pattern along the anteroposterior axis. Their expressions are also complementary in the GPe and GPi: PCDH17 protein is expressed in the inner regions of the GP, whereas PCDH10 protein is expressed in the outer regions of the GP. In the SNr, PCDH17 and PCDH10 proteins are expressed in the posterior and anterior regions, respectively. We also performed *in situ* hybridization experiments to determine the mRNA expression of each *Pcdh* (Fig. 1B) (*10*). We found that the mRNA expression patterns of *Pcdh17* and *Pcdh10* are similar to the protein expression patterns of PCDH17 and PCDH10 in all subregions in the BG, where *Pcdh17* and *Pcdh10* show complementary expression patterns, with a small subset (10-17%) of overlap in each subregion. These results indicate that PCDH17 and PCDH10 are expressed by largely distinct cell populations in the BG subregions and display complementary expression patterns.

**Fig. 1.**
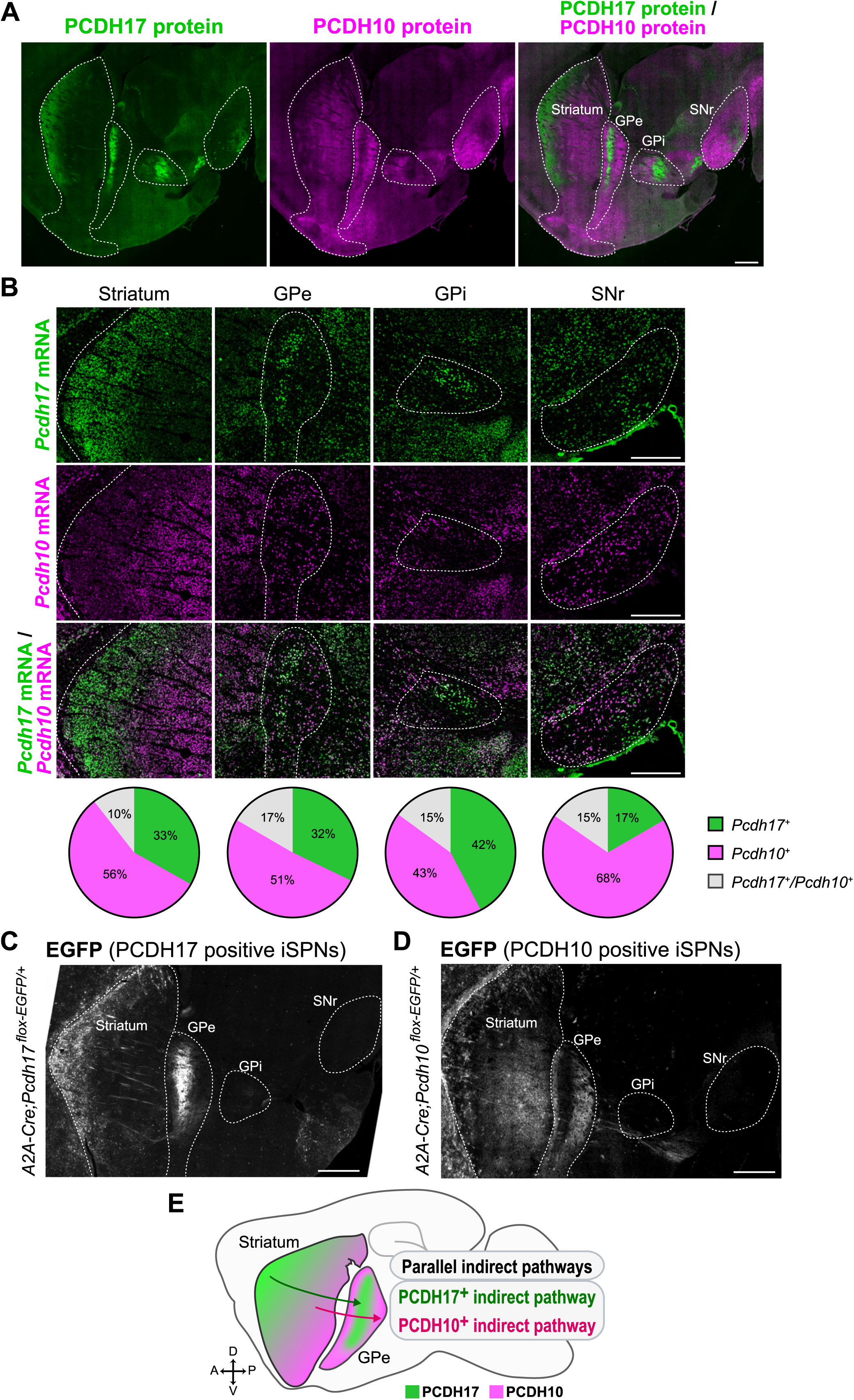
PCDH17 and PCDH10 define parallel iSPN connections in the BG. (**A**) Double staining for PCDH17 (green) and PCDH10 (magenta) proteins in a sagittal BG section from a P8 mouse. PCDH17 and PCDH10 proteins show complementary expression patterns in the BG nuclei. (**B**) Double fluorescent *in situ* hybridization for *Pcdh17* (green) and *Pcdh10* (magenta) mRNAs in a sagittal BG section from a P10 mouse. *Pcdh17* mRNA is preferentially expressed in the anterior striatum and inner GP, while *Pcdh10* mRNA is preferentially expressed in the posterior striatum and outer GP. Reproduced from Hoshina *et al.* (2013) (*10*). Pie charts illustrate the proportions of cells expressing *Pcdh17* only, *Pcdh10* only, and both *Pcdhs* in each basal ganglia nucleus. While most cells only express *Pcdh17* or *Pcdh10*, a subset of cells co-expresses both *Pcdhs*. Number of cells analyzed: 1168 (Striatum), 709 (GPe), 187 (GPi), and 234 (SNr). (**C** to **E**) PCDH17- and PCDH10-expressing iSPNs are connected to PCDH17- and PCDH10-expressing GPe neurons, respectively. EGFP is expressed under the *Pcdh* promoter in iSPNs using *A2A-Cre;Pcdh^flox-EGFP/+^* mice. (**C**) EGFP expression in *A2A-Cre;Pcdh17^flox-EGFP/+^* mice at P10. PCDH17-expressing iSPNs are preferentially localized in the anterior striatum and project to the inner GPe. (**D**) EGFP expression in *A2A-Cre;Pcdh10^flox-EGFP/+^*mice at P12. PCDH10-expressing iSPNs are preferentially localized in the posterior striatum and project to the outer GPe. (**E**) Schematic illustration of the parallel indirect pathways with PCDH17- and PCDH10-expressing iSPNs. The scale bars represent 0.5 mm. GPe, external globus pallidus; GPi, internal globus pallidus; SNr, substantia nigra pars reticulata.

Since δ2-PCDHs are homophilic cell adhesion molecules, we next wanted to know whether BG subregions expressing each PCDH are preferentially connected with each other. Axons from iSPNs in the striatum project to the GPe. To test the idea that the striatal regions expressing PCDH17 and PCDH10 are preferentially connected to the GPe regions expressing PCDH17 and PCDH10, respectively, we generated conditional EGFP knock-in mice (fig. S3). For this, we generated an allele (“flox-EGFP”) in which the first exon of the *Pcdh* gene is flanked by loxP sequences followed by the EGFP expression cassette. EGFP is designed to express under the *Pcdh* promoter, after the first exon of the *Pcdh* gene is removed by Cre recombinase. We mated *Pcdh^flox-EGFP/+^*mice with *A2A-Cre* mice (*11*, *12*), in which Cre is selectively expressed by iSPNs in the striatum, to label *Pcdh*-expressing iSPNs with EGFP. EGFP staining of BG sections from *A2A-Cre;Pcdh17^flox-EGFP/+^* mice showed that *Pcdh17*-expressing iSPNs are distributed in the anterior striatum and their axon terminals project to the inner GPe (Fig. 1C), where *Pcdh17* is highly expressed (Fig. 1, A and B). In *A2A-Cre;Pcdh10^flox-EGFP/+^* mice, *Pcdh10*-expressing iSPNs are distributed in the posterior striatum, and their axon terminals project to the outer GPe (Fig. 1D), where *Pcdh10* is highly expressed (Fig. 1, A and B). EGFP signals were not detected in the GPi and SNr (Fig. 1, C and D), the targets of the direct SPNs (dSPNs). These results suggest that PCDH17- and PCDH10-expressing iSPNs are connected to PCDH17- and PCDH10-expressing GPe neurons, respectively, forming parallel and anatomically segregated indirect pathways (Fig. 1E).

We further examined whether PCDH-expressing iSPNs preferentially form synaptic contacts with GPe neurons expressing the same PCDH. For this, we used *Pcdh17^EGFP/+^* and *Pcdh10^EGFP/+^* knock-in mice, in which EGFP is constitutively expressed under the control of the endogenous *Pcdh* promoter due to germline recombination (fig. S3). In these mice, EGFP labels both PCDH-expressing neurons and their axonal terminals. Inhibitory presynaptic terminals in the GPe, which mainly originate from iSPNs, were identified by immunostaining for VGAT (inhibitory presynaptic marker). PCDH-positive inhibitory terminals were identified as EGFP^+^/VGAT^+^ terminals. We then quantified the number of PCDH-positive inhibitory terminals formed onto PCDH-positive and PCDH-negative GPe neurons. We found that in *Pcdh17^EGFP/+^* mice, PCDH17-positive inhibitory terminals selectively innervated PCDH17-positive GPe neurons, with little innervation on PCDH17-negative neurons (fig. S4, A and B). A similar pattern was observed in *Pcdh10^EGFP/+^* mice, where PCDH10-positive inhibitory terminals preferentially target PCDH10-positive GPe neurons compared to PCDH10-negative GPe neurons. (fig. S4, C and D). These results provide direct anatomical evidence that iSPNs expressing PCDH17 or PCDH10 form synaptic connections with GPe neurons expressing the matching PCDH, supporting the existence of molecularly segregated, parallel indirect pathways.

### Generation of iSPN-specific *Pcdh17* and *Pcdh10* cKO mice

To investigate the roles of PCDH17 and PCDH10 in iSPNs, we generated iSPN-specific *Pcdh* conditional knockout (cKO) mice (fig. S3) by removing both the first exon of the *Pcdh* gene (by mating with *A2A-Cre* mice) and the EGFP cassette (by mating with *ACTB-FLPe* mice). The levels of PCDH17 and PCDH10 proteins were reduced in the striatum and GPe (the target of iSPNs), but not in the SNr (a target of dSPNs), in the cKO mice (fig. S5, A and B). The protein remaining in the striatum would be from dSPNs and input neurons to the striatum. The protein remaining in the GPe would be from GPe neurons (see Fig. 1B). We further examined whether PCDH17 and PCDH10 were selectively deleted in iSPNs and not in dSPNs in the striatum. Since dopamine receptors D1 (D1R) and D2 (D2R) are selectively expressed by dSPNs and iSPNs, respectively (*13*), we performed triple staining for PCDH, D1R, and D2R. In control mice, PCDH17 and PCDH10 signals were detected in the anterior and posterior striatum, respectively, in both iSPNs and dSPNs. In the anterior striatum of *Pcdh17* cKO mice, the PCDH17 signal was diminished in iSPNs but not in dSPNs (fig. S6A). In the posterior striatum of *Pcdh10* cKO, the PCDH10 signal in iSPNs was diminished, while there was no change in dSPNs (fig. S6B). These results indicate that in both *Pcdh17* cKO and *Pcdh10* cKO mice, the expression of PCDH17 and PCDH10 is specifically reduced in iSPNs, but not in dSPNs.

To exclude the possibility that the expression patterns of D1R and D2R are altered in *Pcdh* cKO mice, we labeled iSPNs with tdTomato by crossing *A2A-Cre* mice with *Rosa^STOP-tdT^*mice. The *A2A-Cre* mouse expresses Cre in iSPNs independent of D2R expression (*11*, *12*), allowing us to identify iSPNs as tdTomato⁺ cells. Using this strategy, we found that in *Pcdh17* control (tdT), *Pcdh17* cKO (tdT), *Pcdh10* control (tdT), and *Pcdh10* cKO (tdT) mice, tdTomato⁺ cells are D2R-positive (fig. S6, C and D), indicating that the expression pattern of dopamine receptors is not changed in *Pcdh* cKO mice.

To determine whether the loss of one PCDH protein affects the expression pattern of the other, we analyzed PCDH10 protein expression in iSPN-*Pcdh17* cKO mice, and PCDH17 protein expression in iSPN-*Pcdh10* cKO mice. In *Pcdh17* cKO mice, the PCDH10 expression pattern and levels were not changed in the anterior striatum and inner GPe, where PCDH17 levels were decreased (fig. S5C). Similarly, in *Pcdh10* cKO mice, the PCDH17 expression pattern and levels were not changed in the posterior striatum and outer GPe, where PCDH10 levels were decreased (fig. S5D). These findings suggest that the loss of PCDH17 does not affect PCDH10 expression, and vice versa.

### Region-specific abnormal SPN aggregations in iSPN-*Pcdh* cKO mice

The *iSPN-Pcdh17* cKO and *iSPN-Pcdh10 cKO* mice did not die early, and the mice were fertile. In addition, the gross appearance of the brains of both cKO mice was normal. Using the iSPN-*Pcdh* cKO mice, we investigated the SPN distribution in the striatum, axonal targeting by iSPNs, SPN dendrite morphology, synapses formed by iSPNs, synapses formed onto iSPNs, synaptic physiology, and the behavioral consequences of *Pcdh* deletion.

We first examined the distribution of dSPNs and iSPNs in the striatum of iSPN-*Pcdh* cKO mice by staining for D1R and D2R at 3–4 weeks of age. In the normal striatum at this age, SPNs expressing D1R and D2R are evenly distributed (*14*). In *Pcdh17* cKO mice, we found that D1R and D2R signals showed abnormal aggregations in the anterior striatum, where PCDH17 is highly expressed (Fig. 2, A and B). D1R and D2R signals were evenly distributed in the posterior striatum of *Pcdh17* cKO mice. In contrast, in *Pcdh10* cKO mice, D1R and D2R signals showed abnormal aggregations only in the posterior striatum, where PCDH10 is highly expressed (Fig. 2, C and D).

**Fig. 2.**
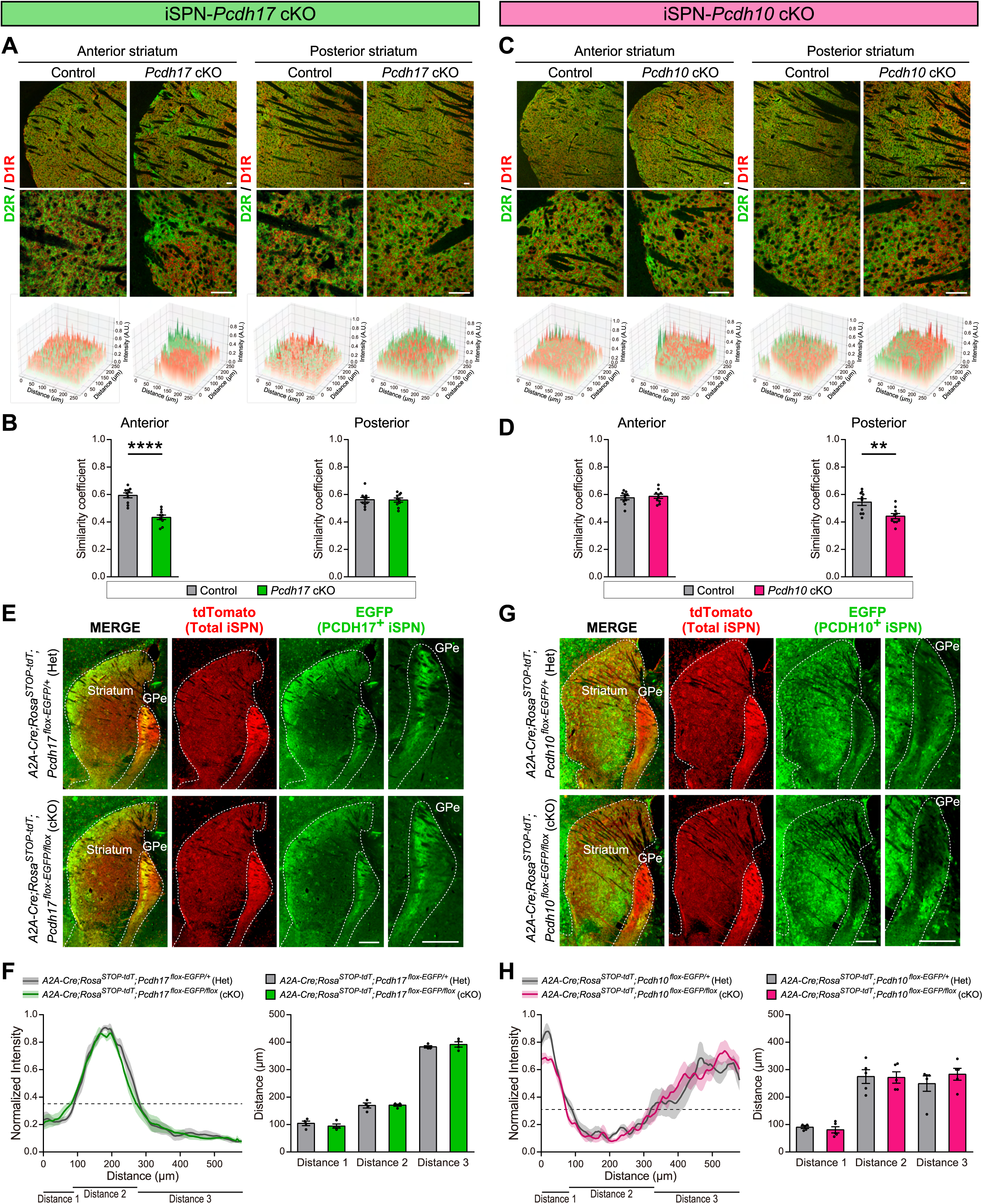
Abnormal aggregation of SPNs in the striatum and normal iSPN axonal targeting in iSPN-*Pcdh* cKO mice. (**A** to **D**) Region-specific abnormal sorting of dSPNs and iSPNs in iSPN-*Pcdh* cKO mice. (**A**) Double staining for D2R (green; labels iSPNs) and D1R (red; labels dSPNs) of anterior and posterior striatal sections from 3- to 4-week-old *Pcdh17* cKO and control mice. The upper panels show low magnification images, and the lower panels show high magnification images. Contour maps for D2R (green) and D1R (red) in the anterior and posterior striatum are shown below. (**B**) Quantification of the similarity coefficient in the anterior and posterior striatum. *n* = 10 (total) fields from 2 mice per genotype. (**C**) Double staining for D2R and D1R of anterior and posterior striatal sections from 3- to 4-week-old *Pcdh10* cKO and control mice, and contour maps. (**D**) Quantification of the similarity coefficient in the anterior and posterior striatum. *n* = 10 (total) fields from 2 mice per genotype. (**E** to **H**) Normal iSPN axonal targeting in iSPN-*Pcdh* cKO (*A2A-Cre;Rosa^STOP-tdT^;Pcdh^flox-EGFP/flox^*) mice. (**E**) Immunostaining for tdTomato (red: labels all iSPNs) and EGFP (green: labels PCDH17-expressing iSPNs) of BG sections from adult *A2A-Cre;Rosa^STOP-tdT^;Pcdh17^flox-EGFP/+^*(Het) and *A2A-Cre;Rosa^STOP-tdT^;Pcdh17^flox-EGFP/flox^*(cKO) mice. (**F**) Quantification of the EGFP intensities in the GPe. Line scan (left; dotted line indicates background intensity) and quantification of distances (right) are shown in the graphs. See fig. S9 for the method. The axonal targeting of PCDH17-expressing iSPNs to the GPe is similar between Het and cKO mice. *n* = 4 mice per genotype. (**G**) Immunostaining for tdTomato (red) and EGFP (green: PCDH10-expressing iSPNs are labeled) of BG sections from adult *A2A-Cre;Rosa^STOP-tdT^;Pcdh10^flox-EGFP/+^*(Het) and *A2A-Cre;Rosa^STOP-tdT^;Pcdh10^flox-EGFP/flox^*(cKO) mice. (**H**) Quantification of the EGFP intensities in the GPe. The axonal targeting of PCDH10-expressing iSPNs to the GPe is similar between Het and cKO mice. *n* = 5 mice per genotype. The scale bars represent 50 μm (**A** and **C**) and 0.5 mm (**E** and **G**). Data are mean ± SEM. \*\**P* < 0.01, \*\*\*\**P* < 0.0001 by Student’s *t*-test.

The changes in D1R/D2R signals could be due to changes in SPN distribution and/or changes in SPN morphology. To further assess SPN distribution, we identified the cell bodies of dSPNs and iSPNs based on D1R and D2R expression (fig. S7, A and C; also see fig. S6) (*13*). We then performed nearest neighbor analysis to quantify the clustering of dSPNs and iSPNs in *Pcdh17* cKO and *Pcdh10* cKO mice. We found that in the anterior striatum, but not the posterior striatum, of *Pcdh17* cKO mice the nearest neighbor distances for dSPNs and iSPNs were reduced relative to control mice (fig. S7, A and B). On the other hand, in the posterior striatum, but not the anterior striatum, of *Pcdh10* cKO mice, the nearest neighbor distances for dSPNs and iSPNs were reduced relative to control mice (fig. S7, C and D). These results indicate that in regions where each PCDH is abundant, PCDH-positive dSPNs and PCDH-negative iSPNs were segregated and abnormally clustered, suggesting that PCDH17 and PCDH10 play important roles in evenly distributing dSPNs and iSPNs in a region-specific manner.

To assess SPN morphology, we evaluated cell body size and dendritic morphology by sparsely labeling SPNs. For this, we retro-orbitally injected adeno-associated viruses (AAVs) expressing membrane-tagged EGFP into *Pcdh17* cKO (tdT) and *Pcdh10* cKO (tdT) mice, in which iSPNs are labeled with tdTomato. tdTomato⁺/mEGFP⁺ striatal cells were identified as iSPNs, while tdTomato⁻/mEGFP⁺ striatal cells were identified as dSPNs. We found that the size of cell bodies and the complexity of dendrites, assessed by the number of dendritic crossings, were normal in both the anterior and posterior striatum of *Pcdh17* cKO (tdT) and *Pcdh10* cKO (tdT) mice (fig. S8, A to D). These findings suggest that the loss of PCDH17 or PCDH10 in iSPNs does not affect the morphology of either dSPNs or iSPNs.

To determine when abnormal SPN aggregations occur during development, we examined the distribution of dSPNs and iSPNs at an earlier time point, postnatal day 0 (P0). We found no differences in the distribution patterns of dSPNs and iSPNs at P0 in either the anterior or posterior striatum of *Pcdh17* cKO and *Pcdh10* cKO mice (fig. S7, E to H). Thus, while the distribution patterns of dSPNs and iSPNs are similar immediately after birth, differences emerge by 3-4 weeks of age. The striatum contains two compartments termed the patch and the matrix. In wild-type mice at P0, both dSPNs and iSPNs are highly enriched in the patch structure of the striatum. By P28, they are distributed into both the patch and matrix structures (*15*). Therefore, abnormal aggregations occur when SPNs distribute to the patch and matrix structures between P0 and P28 in iSPN-*Pcdh* cKO mice.

### Normal iSPN axonal targeting in iSPN-*Pcdh* cKO mice

We next asked whether PCDH17 and PCDH10 regulate the axonal projections of iSPNs. To visualize *Pcdh*-expressing iSPN axons, we utilized *A2A-Cre;Rosa^STOP-tdT^;Pcdh^flox-EGFP^* mice, in which all iSPNs express tdTomato, and *Pcdh*-expressing iSPNs express EGFP (Fig. 2, E and G). We compared iSPN axonal projections between adult *A2A-Cre; Rosa^STOP-tdT^; Pcdh^flox-EGFP/+^* (Het) and *A2A-Cre;Rosa^STOP-tdT^;Pcdh^flox-EGFP/flox^ (cKO)* mice. We assessed the targeting of PCDH17-and PCDH10-expressing iSPN axons in the GPe by evaluating the EGFP-positive regions in the GPe (Fig. 2, E to H; see fig. S9, A to G for the evaluation method). We did not find differences in the EGFP expression patterns between Het and cKO mice. The density of iSPN axon terminals in the GPe, assessed by quantifying the EGFP intensity in the GPe (normalized to that in the striatum), was also similar between Het and cKO mice for both PCDHs (fig. S9, H and I). These results suggest that the targeting and density of iSPN axon terminals in the GPe are normal in the absence of PCDH17 or PCDH10.

### Synaptic defects in iSPN-*Pcdh* cKO mice

PCDH17 and PCDH10 are localized at both excitatory and inhibitory synapses in the mouse brain (*10*, *16*). Therefore, we investigated iSPN synapses. We first investigated synapses formed by the iSPNs, i.e., inhibitory synapses in the GPe. For this, we quantified the densities and sizes of VGAT (inhibitory presynaptic marker) and Gephyrin (inhibitory postsynaptic marker) puncta in the inner GPe, the target of PCDH17-expressing iSPNs (Fig. 1C), and the outer GPe, the target of PCDH10-expressing iSPNs (Fig. 1D). In *Pcdh17* cKO mice, the density of VGAT was significantly decreased in the inner GPe (Fig. 3A; *P* = 0.0003; Mean Difference [MD] = -11.7), but not in the outer GPe (Fig. 3B). The size of VGAT puncta was increased in the outer GPe (Fig. 3B). In contrast, in *Pcdh10* cKO mice, the density of VGAT was significantly decreased in the outer GPe (Fig. 3D; *P* = 0.0002; MD = -17.7), but not in the inner GPe (Fig. 3C). The size of VGAT puncta was similar between *Pcdh10* cKO and control mice (Fig. 3, C and D). To compare the puncta density of VGAT between the inner GPe and outer GPe, we normalized the puncta density in *Pcdh* cKO mice to that in relevant controls and compared the normalized density between the two regions. We found a significant decrease in VGAT density in the inner GPe relative to outer GPe in *Pcdh17* cKO mice (fig. S10A; *P* = 0.0178). A modest reduction was also observed in the outer GPe compared to the inner GPe in *Pcdh10* cKO mice (*P* = 0.0764; fig. S10B).

**Fig. 3.**
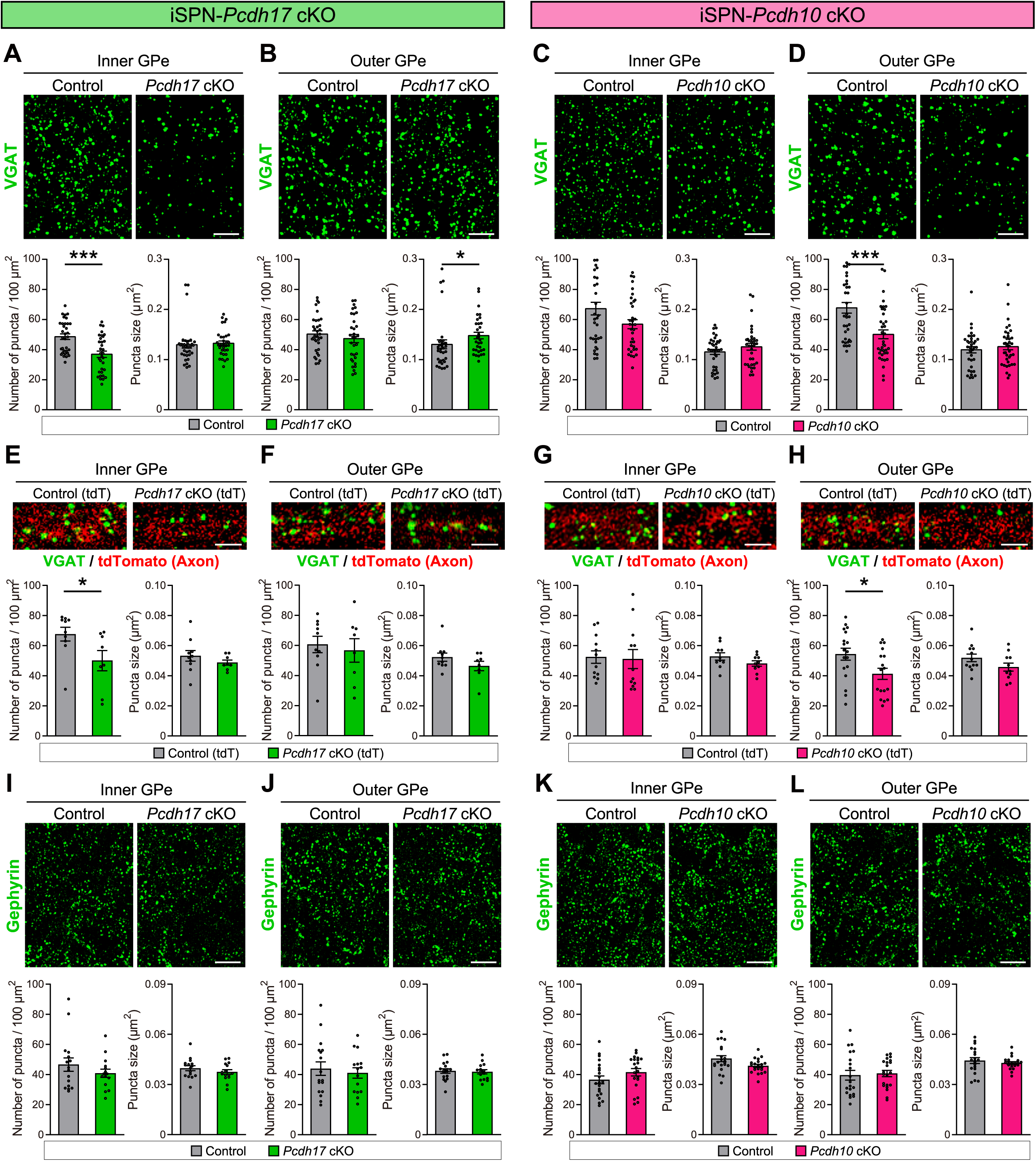
Impaired inhibitory synaptogenesis by iSPNs in the GPe of *Pcdh17* cKO and *Pcdh10* cKO mice. VGAT puncta (**A** to **D**), VGAT puncta in iSPN axons (**E** to **H**), and Gephyrin puncta (**I** to **L**) in the inner GPe (iGPe) and outer GPe (oGPe) of 2-week-old *Pcdh* cKO and control mice. (**A** and **B**) The density of VGAT puncta in the iGPe (**A**; *P* = 0.0003), but not oGPe (**B**), is decreased in *Pcdh17* cKO mice relative to control mice. The size of VGAT puncta in oGPe (**B**; *P* = 0.0101), but not iGPe (**A**), is increased in *Pcdh17* cKO mice relative to control mice. *n* = 32-33 fields from 8 mice per genotype. (**C** and **D**) The density of VGAT puncta in the oGPe (**D**; *P* = 0.0002), but not iGPe (**C**), is decreased in *Pcdh10* cKO mice relative to control mice. The size of VGAT puncta in the iGPe and oGPe is similar between *Pcdh10* cKO and control mice. *n* = 35-37 fields from 10 mice per genotype. (**E** and **F**) The density of VGAT puncta in iSPN axons in the iGPe (**E**; *P* = 0.0250), but not oGPe (**F**), is decreased in *Pcdh17* cKO (tdT) relative to control (tdT) mice. The size of VGAT puncta in iSPN axons in the iGPe and oGPe is similar between *Pcdh17* cKO (tdT) and control (tdT) mice. *n* = 8-10 fields from 4 mice per genotype. (**G** and **H**) The density of VGAT puncta in iSPN axons in the oGPe (**H**; *P* = 0.0386), but not iGPe (**G**), is decreased in *Pcdh10* cKO (tdT) relative to control (tdT) mice. The size of VGAT puncta in iSPN axons in the iGPe and oGPe is similar between *Pcdh10* cKO (tdT) and control (tdT) mice. Density; *n* = 12-18 fields from 6 mice per genotype. Size; *n* = 10-12 fields from 4 mice per genotype. (**I** and **J**) The density and size of Gephyrin puncta in the iGPe (**I**) and oGPe (**J**) are similar between *Pcdh17* cKO and control mice. *n* = 15-17 fields from 4 mice per genotype. (**K** and **L**) The density and size of Gephyrin puncta in the iGPe (**K**) and oGPe (**L**) are similar between *Pcdh10* cKO and control mice. *n* = 20-21 fields from 5 mice per genotype. The scale bars represent 2 μm. Data are mean ± SEM. \**P* < 0.05, \*\**P* < 0.01, \*\*\**P* < 0.001, \*\*\*\**P* < 0.0001 by Mann-Whitney U test.

To specifically analyze VGAT signals in iSPN axons, we labeled iSPN axons with tdTomato by crossing *Pcdh* cKO mice with *Rosa^STOP-tdT^*mice (tdT). In *Pcdh17* cKO (tdT) mice, VGAT density was decreased in iSPN axons in the inner GPe (Fig. 3E; *P* = 0.0250), but not in the outer GPe (Fig. 3F). In contrast, in *Pcdh10* cKO (tdT) mice, VGAT density was decreased in iSPN axons in the outer GPe (Fig. 3H; *P* = 0.0386), but not in the inner GPe (Fig. 3G). The size of VGAT puncta in iSPN axons did not differ between *Pcdh* cKO (tdT) and control (tdT) mice (Fig. 3, E to H). To functionally assess inhibitory synapses, we performed electrophysiological recordings of spontaneous inhibitory postsynaptic currents (sIPSCs) from neurons in the outer GPe. In *Pcdh17* cKO (tdT) mice, both the amplitude and frequency of sIPSCs were comparable to control (tdT) mice (fig. S11A). However, in *Pcdh10* cKO (tdT) mice, the frequency of sIPSCs was significantly reduced in the outer GPe compared to control (tdT) mice, while the amplitude remained unchanged (fig. S11B), suggesting an impaired inhibitory presynaptic function in the outer GPe of *Pcdh10* cKO (tdT) mice. The density and size of Gephyrin puncta were similar between *Pcdh* cKO and control mice (Fig. 3, I to L). Together, these results indicate that in *Pcdh17* cKO and *Pcdh10* cKO mice, inhibitory presynaptic development is preferentially impaired in the region of the GPe targeted by PCDH17- or PCDH10-expressing iSPNs.

We next investigated excitatory synapses formed onto iSPNs in the striatum of iSPN-*Pcdh* cKO mice. Striatal iSPNs receive VGLUT1-positive excitatory synapses from the cerebral cortex and VGLUT2-positive excitatory synapses from the thalamus (*17*). We have previously shown that PCDH17 and PCDH10 show complementary expressions in the cortex and thalamus as well (*10*). Therefore, we examined these types of excitatory synapses on iSPNs. To evaluate presynaptic development, we examined the densities and sizes of VGLUT1 and VGLUT2 puncta on iSPN dendrites in the anterior and posterior striatum. iSPN dendrites were labeled with tdTomato by mating with *Rosa^STOP-tdT^* mice (tdT). The density of VGLUT1 puncta on iSPN dendrites was decreased both in the anterior (Fig. 4A; *P* < 0.0001; MD = -9.99) and posterior (Fig. 4B; *P* = 0.0023; MD = -5.31) striatum of *Pcdh17* cKO (tdT) mice compared to control (tdT) mice. The size of VGLUT1 puncta was only decreased in the anterior striatum (Fig. 4, A and B). In *Pcdh10* cKO (tdT) mice, the density of VGLUT1 puncta was also reduced in the anterior (Fig. 4C; *P* = 0.0315; MD = -10.0) and posterior (Fig. 4D; *P* = 0.0080; MD = -15.6) striatum. We then compared the normalized puncta density of VGLUT1 between the anterior and posterior striatum. We found a significant decrease in VGLUT1 density in the anterior striatum relative to the posterior striatum in *Pcdh17* cKO (tdT) mice (fig. S10C; *P* = 0.0371), while it was similar between the anterior and posterior regions in *Pcdh10* cKO (tdT) mice (fig. S10D; *P* = 0.3427).

**Fig. 4.**
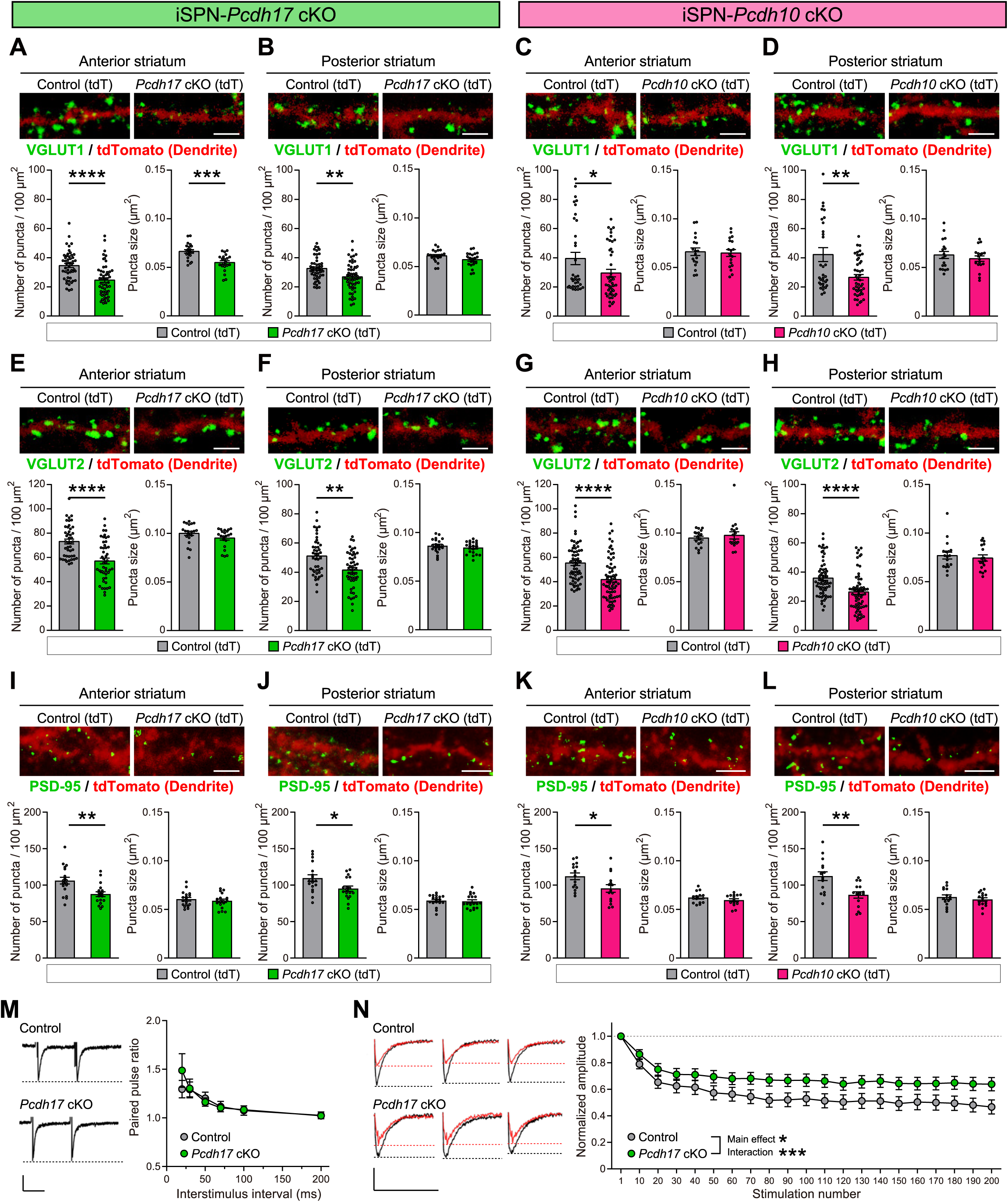
Reduced excitatory synapses formed on iSPNs *in Pcdh17* cKO and *Pcdh10* cKO mice. (**A** to **L**) VGLUT1 (**A** to **D**), VGLUT2 (**E** to **H**), and PSD-95 (**I** to **L**) puncta on iSPN dendrites (labeled with tdT) in the anterior and posterior striatum of 2-week-old *Pcdh* cKO and control mice. (**A** and **B**) The density of VGLUT1 puncta on iSPNs in both the anterior (**A;** *P* < 0.0001) and posterior (**B;** *P* = 0.0023) striatum is decreased in *Pcdh17* cKO (tdT) mice relative to control mice (tdT). The size of VGLUT1 puncta on iSPNs is only decreased in the anterior striatum (**A**; *P* = 0.0001). Density; *n* = 48-54 fields from 7 mice per genotype. Size; *n* = 21 fields from 7 mice per genotype. (**C** and **D**) The density of VGLUT1 puncta on iSPNs in both the anterior (**C;** *P* = 0.0315) and posterior (**D;** *P* = 0.0080) striatum is decreased in *Pcdh10* cKO (tdT) mice relative to control (tdT) mice. Density; *n* = 33-45 fields from 6 mice per genotype. Size; *n* = 18 fields from 6 mice per genotype. (**E** and **F**) The density of VGLUT2 puncta on iSPNs in both the anterior (**E;** *P* < 0.0001) and posterior (**F;** *P* = 0.0014) striatum is decreased in *Pcdh17* cKO (tdT) mice relative to control (tdT) mice. Density; *n* = 44-50 fields from 7 mice per genotype. Size; *n* = 20-21 fields from 7 mice per genotype. (**G** and **H**) The density of VGLUT2 puncta on iSPNs in both the anterior (**G**; *P* < 0.0001) and posterior (**H**; *P* < 0.0001) striatum is decreased in *Pcdh10* cKO (tdT) mice relative to control (tdT) mice. Density; *n* = 63-68 fields from 10 mice per genotype. Size; *n* = 17-18 fields from 6 mice per genotype. (**I** and **J**) The density of PSD-95 puncta on iSPNs in both the anterior (**I;** *P* = 0.0037) and posterior (**J;** *P* = 0.0290) striatum is decreased in *Pcdh17* cKO (tdT) mice relative to control (tdT) mice. *n* = 18 fields from 6 mice per genotype. (**K** and **L**) The density of PSD-95 puncta on iSPNs in both the anterior (**K;** *P* = 0.0367) and posterior (**L;** *P* = 0.0043) striatum is decreased in *Pcdh10* cKO (tdT) mice relative to control (tdT) mice. *n* = 15 fields from 5 mice per genotype. **(M)** Paired-pulse ratio (PPR) of evoked EPSCs at cortico-striatal synapses in the anterior striatum of P11–P23 *Pcdh17* cKO and control mice. (Left) Sample traces with a 20-ms interstimulus interval. (Right) PPR across a range of interstimulus intervals (20, 30, 50, 70, 100, and 200 ms). PPR is similar between *Pcdh17* cKO and control mice. *n* = 35 neurons from 6 control mice; 39 neurons from 6 *Pcdh17 cKO* mice. **(N)** Synaptic depression induced by prolonged repetitive stimulation (10 Hz, 200 stimuli) at cortico-striatal synapses in the anterior striatum of P11–P23 *Pcdh17* cKO and control mice. (Left) Sample traces showing responses 1–3 (black) and 198–200 (red). (Right) Quantification of normalized EPSC amplitudes (each point represents the average of 10 consecutive responses) reveals weaker synaptic depression in *Pcdh17* cKO mice compared to controls (main effect of genotype: *P* = 0.0493; genotype × stimulus interaction: *P* = 0.0002). *n* = 39 neurons from 6 control mice; 45 neurons from 6 *Pcdh17* cKO mice. The scale bars represent 2 μm (**A** to **L**) and 100 pA and 10 ms (**M** and **N**). Data are mean ± SEM. \**P* < 0.05, \*\**P* < 0.01, \*\*\**P* < 0.001, \*\*\*\**P* < 0.0001 by Mann-Whitney U test (**A** to **L**), two-way ANOVA followed by the Sidak test (**M**) and two-way repeated-measures ANOVA followed by the Sidak test (**N**).

The density of VGLUT2 puncta was reduced in both the anterior (Fig. 4E; *P* < 0.0001; MD = -16.0) and posterior (Fig. 4F; *P* = 0.0014; MD = -9.67) striatum in *Pcdh17* cKO (tdT) mice. In *Pcdh10* cKO (tdT) mice, the density of VGLUT2 puncta was also reduced in the anterior (Fig. 4G; *P* < 0.0001; MD = -13.5) and posterior (Fig. 4H; *P* < 0.0001; MD = -9.56) striatum. The normalized puncta density of VGLUT2 was comparable between the anterior and posterior regions in *Pcdh17* and *Pcdh10* cKO (tdT) mice (fig. S10, E and F; *P* = 0.3900 and *P* = 0.5756).

To evaluate postsynaptic development, we examined the clustering of PSD-95 along the iSPN dendrites. We found that the density of PSD-95 puncta was decreased both in the anterior (Fig. 4I; *P* = 0.0037 for *Pcdh17* cKO and 0.0367 for *Pcdh10* cKO) and posterior (Fig. 4J; *P* = 0.0290 for *Pcdh17* cKO and 0.0043 for *Pcdh10* cKO) striatum of *Pcdh* cKO (tdT) mice. These results indicate that *Pcdh17* cKO and *Pcdh10* cKO mice show impaired development of excitatory synapses formed onto the iSPNs. The results suggest that there appears to be more robust decreases in the areas where each PCDH is highly expressed (e.g., VGLUT1 in *Pcdh17* cKO mice), but also show that changes were observed outside of the regions with high expression, which may reflect complex feedback in the circuit or be secondary to abnormal cell aggregation.

We next analyzed short-term synaptic plasticity at cortico-striatal excitatory synapses in *Pcdh17* cKO mice, as global *Pcdh17* KO mice have been shown to exhibit altered short-term synaptic plasticity (*10*). We found that, while the paired-pulse ratio (PPR) was not significantly different between control and *Pcdh17* cKO mice (Fig. 4M), synaptic depression by prolonged repetitive stimulation was significantly weaker in *Pcdh17* cKO mice compared to controls (Fig. 4N; main effect, *P* = 0.0493; interaction, *P* = 0.0002). These results suggest that, like global *Pcdh17* KO mice, *Pcdh17* cKO mice show impaired presynaptic short-term plasticity.

We then examined inhibitory synapses on iSPNs. For this, we quantified the densities and sizes of VGAT and Gephyrin puncta on iSPN cell bodies, because many inhibitory synapses are formed around iSPN cell bodies. We found that the density of VGAT puncta on iSPN cell bodies was decreased both in the anterior (Fig. 5A; *P* = 0.0035; MD = -2.06) and posterior (Fig. 5B; *P* = 0.0138; MD = -1.91) striatum of *Pcdh17* cKO (tdT) mice. In *Pcdh10* cKO (tdT) mice, the density of VGAT puncta was also decreased in the posterior striatum (Fig. 5D; *P* < 0.0001; MD = -3.64) and the anterior striatum (Fig. 5C; *P* = 0.0005; MD = -3.18). The normalized puncta density of VGAT on iSPNs was comparable between the anterior and posterior regions in *Pcdh17* cKO (tdT) mice (fig. S10G; *P* = 0.7646), while a modest reduction was observed in the posterior regions compared to the anterior regions in *Pcdh10* cKO (tdT) mice (fig. S10H; *P* = 0.0963). The density and size of Gephyrin puncta were similar in *Pcdh17* cKO and *Pcdh10* cKO mice compared to their control mice (Fig. 5, E to H). These results suggest that *Pcdh17* cKO and *Pcdh10* cKO mice show impaired presynaptic development of inhibitory synapses on iSPNs.

**Fig. 5.**
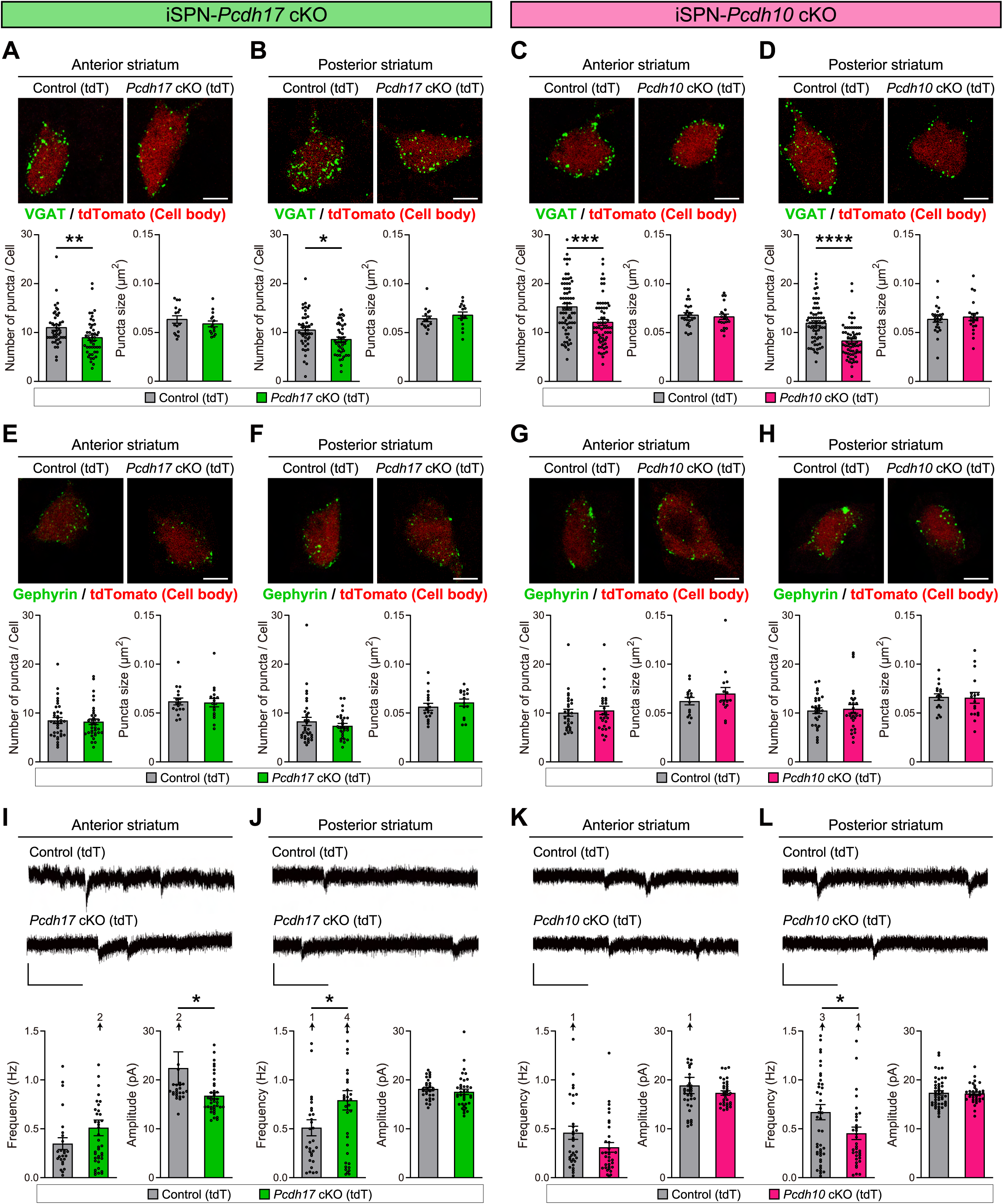
*Pcdh17* cKO and *Pcdh10* cKO mice exhibit defects in inhibitory synapses on iSPNs and impaired excitatory synaptic function in iSPNs. (**A** to **H**) VGAT (**A** to **D**) and Gephyrin (**E** to **H**) puncta on iSPN cell bodies (labeled with tdT) in the anterior and posterior striatum of 2-week-old *Pcdh* cKO and control mice. (**A** and **B**) The density of VGAT puncta on iSPN cell bodies in both the anterior (**A;** *P* = 0.0035) and posterior (**B;** *P* = 0.0138) striatum is decreased in *Pcdh17* cKO (tdT) mice relative to control (tdT) mice. Density; *n* = 43-47 fields from 7 mice per genotype. Size; *n* = 15 fields from 5 mice per genotype. (**C** and **D**) The density of VGAT puncta on iSPN cell bodies in both the anterior (**C;** *P* = 0.0005) and posterior (**D;** *P* < 0.0001) striatum is decreased in *Pcdh10* cKO (tdT) mice relative to control (tdT) mice. Density; *n* = 62-70 fields from 6 mice per genotype. Size; *n* = 21-24 fields from 9 mice per genotype. (**E** and **F**) The density and size of Gephyrin puncta on iSPN cell bodies in the anterior (**E**) and posterior (**F**) striatum are similar between *Pcdh17* cKO (tdT) and control (tdT) mice. Density; *n* = 28-36 fields from 6 mice per genotype. Size; *n* = 15-18 fields from 5 mice per genotype. (**G** and **H**) The density and size of Gephyrin puncta on iSPN cell bodies in the anterior (**G**) and posterior (**H**) striatum are similar between *Pcdh10* cKO (tdT) and control (tdT) mice. Density; *n* = 27-31 fields from 4 mice per genotype. Size; *n* = 15-17 fields from 4 mice per genotype. (**I** to **L**) Spontaneous EPSCs (sEPSCs) recorded from iSPNs in the anterior and posterior striatum. Top panels show example traces. The graphs show the frequencies and amplitudes of sEPSCs in *Pcdh* cKO and control mice. *n* = 25 (anterior) and 30 (posterior) neurons from 7 *Pcdh17* control (tdT) mice; 40 *(anterior) and 37* (posterior) neurons from 8 *Pcdh17 cKO* (tdT) mice; 34 (anterior) and 46 (posterior) neurons from 5 *Pcdh10* control (tdT) mice; and 33 (anterior) and 35 (posterior) neurons from 5 *Pcdh10 cKO* (tdT) mice. The scale bars represent 2 μm (**A** to **H**) and 20 pA and 100 ms (**I** to **L**). Data are mean ± SEM. \**P* < 0.05, \*\**P* < 0.01, \*\*\**P* < 0.001, \*\*\*\**P* < 0.0001 by Mann-Whitney U test (**A** to **H**) and Student’s *t*-test (**I** to **L**).

To investigate the functional consequences of the synaptic defects in *Pcdh17* cKO and *Pcdh10* cKO mice, we recorded spontaneous excitatory postsynaptic currents (sEPSCs) from iSPNs (labeled with tdT) in the anterior and posterior striatum. In *Pcdh17* cKO (tdT) mice, the amplitude of sEPSCs was significantly reduced in the anterior striatum (Fig. 5I; *P* = 0.0431), but not in the posterior striatum (Fig. 5J). The frequency of sEPSCs was not changed in the anterior striatum in *Pcdh17* cKO (tdT) mice; however, it was increased in the posterior striatum. The EPSC frequency can be affected by various factors, including the number of synapses and neurotransmitter release probability. In the posterior striatum of *Pcdh17* cKO mice, compensatory presynaptic changes may have increased neurotransmitter release probability or vesicle release efficiency, thereby increasing the EPSC frequency despite a reduction in synapse number. In *Pcdh10* cKO (tdT) mice, the frequency of sEPSCs was significantly decreased in the posterior striatum (Fig. 5L; *P =* 0.0422), but not in the anterior striatum (Fig. 5K). The amplitude of sEPSC was not changed in *Pcdh10* cKO (tdT) mice. These results indicate that synaptic currents from iSPNs are altered both in *Pcdh17* cKO and *Pcdh10* cKO mice, but the changes are different between them. The complex physiological changes may be because PCDH17 and PCDH10 are involved in both excitatory and inhibitory synapse development (Fig. 4 and Fig. 5). It is also possible that abnormal aggregation of iSPNs (Fig. 2, A to D) is affecting their physiology. Additionally, iSPNs in the anterior and posterior striatum may be influencing each other.

### Impaired task learning and behavioral flexibility in *Pcdh17* cKO mice, but not in *Pcdh10* **cKO mice**

Given the differentially impaired synaptic development of the indirect BG circuits in *Pcdh17* cKO and *Pcdh10* cKO mice, we next investigated the behavioral consequences of *Pcdh* inactivation to determine what kind of behavior is regulated by the circuits organized by PCDH17 and PCDH10. For this, we performed several behavioral paradigms associated with BG functions, including task learning (*18*), motor and sensory behaviors (*19*), and anxiety-related behaviors (*20*).

To evaluate task learning, we used an operant learning system, a reward-based learning system that has been used to assess many types of goal-directed learning, cognitive, and executive functions (*21*, *22*). The mice first went through three stages of pre-training: acclimation to the operant chamber, reward acquisition training, and stimulus touch training (fig. S12, A to F). During pre-training, days to criterion and reaction times (reward collection latency and touch latency) were similar between iSPN-*Pcdh* cKO and control mice, indicating that neither cKO mice have abnormalities in simple reward-based behaviors. Mice that met the criterion in pre-training then proceeded to the pairwise visual discrimination (perception-based cognitive learning) task. In this task, mice were rewarded when they touched the correct image from a pair of visual images (Fig. 6A; Visual discrimination task). Mice performed 60 trials (= one session) per day (Fig. 6B). We found that *Pcdh17* cKO mice acquired the task more slowly than controls (Fig. 6C; main effect, *P* = 0.0489; interaction, *P* = 0.2914). They required more total trials and made more total errors before reaching the learning criterion (average correct response of 80% for 2 consecutive days) compared to control mice (Fig. 6E). On the other hand, *Pcdh10* cKO mice are comparable to control mice in terms of the percentage of correct responses across sessions (Fig. 6D) and the total number of trials and total errors to reach the criterion (Fig. 6F). Neither *Pcdh17* cKO nor *Pcdh10* cKO mice showed any change in the correct response latency compared to their controls (fig. S12, G and H), suggesting that the response speed is normal in both *Pcdh cKO* mice. Together, *Pcdh17* cKO mice, but not *Pcdh10* cKO mice, show impaired perception-based cognitive learning.

**Fig. 6.**
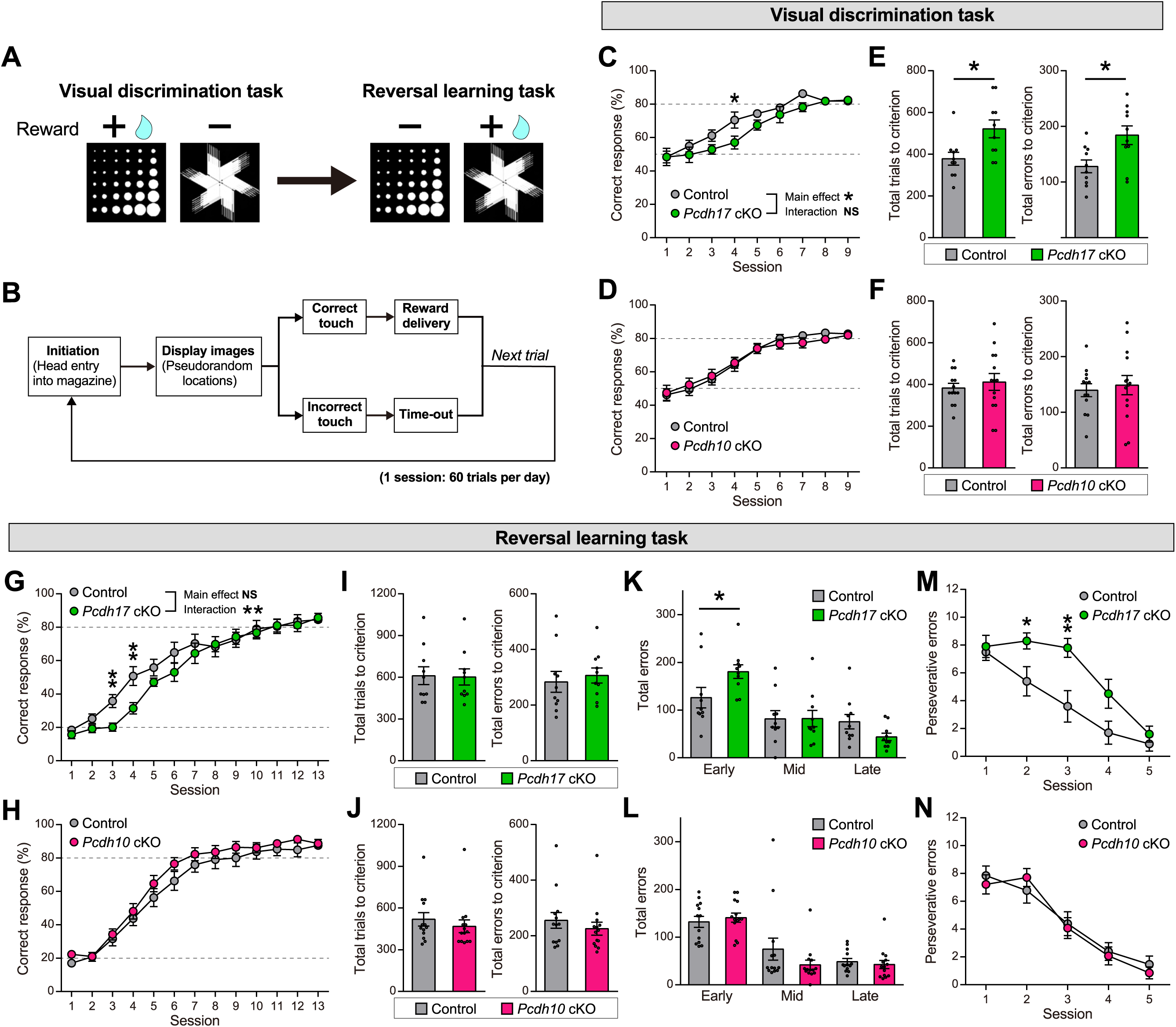
*Pcdh17* cKO but not *Pcdh10* cKO mice show impaired task learning. (A) Schematic of the visual discrimination task (left) and reversal learning task (right). Rewarded stimulus (+) and unrewarded stimulus (−) are represented. In the reversal learning task, + and − images are reversed from the visual discrimination task. (B) Timeline of one trial in the task. One session ends after 60 trials per day. (**C** to **F**) Visual discrimination task. (**C** and **D**) Correct response (%) is decreased in *Pcdh17* cKO mice relative to control mice in session 4. Two-way repeated-measures ANOVA showed a significant main effect of genotype (*P* = 0.0489), but no significant genotype × session interaction (*P* = 0.2914) (**C**). Correct responses are similar in *Pcdh10* cKO mice relative to control mice (**D**). (**E** and **F**) Total trials required to criterion and total errors made to criterion are increased in *Pcdh17* cKO mice (**E**) but not in *Pcdh10* cKO mice (**F**) relative to control mice. *n* = 10 mice for *Pcdh17 control*; 10 for *Pcdh17 cKO;* 13 for *Pcdh10 control*; and 14 for *Pcdh10 cKO*. (**G** to **N**) Reversal learning task. (**G** and **H**) Correct responses are reduced in *Pcdh17* cKO mice relative to control mice in sessions 3 and 4 (**G**). Two-way repeated-measures ANOVA revealed a significant genotype × session interaction (*P* = 0.0044), but no significant main effect of genotype (*P* = 0.2346). Correct responses are similar in *Pcdh10* cKO mice relative to control mice (**H**). (**I** and **J**) Total trials required to criterion and total errors made to criterion are similar in *Pcdh17* cKO mice (**I**) and in *Pcdh10* cKO mice (**J**) relative to control mice. (**K** and **L**) Total errors in the early (0–40% correct response), mid (40–60% correct response), and late (60–80% correct response) phases. Total errors are increased in the early phase in *Pcdh17* cKO mice relative to control mice (**K**). Total errors in all phases are similar in *Pcdh10* cKO mice relative to control mice (**L**). (**M** and **N**) Perseverative errors (four consecutive errors) during the reversal learning sessions 1 to 5. *Pcdh17* cKO mice made more perseverative errors than control mice in sessions 2 and 3. *n* = 10 mice for *Pcdh17 control*; 10 for *Pcdh17 cKO;* 13 for *Pcdh10 control*; and 14 for *Pcdh10 cKO*. Data are mean ± SEM. \**P* < 0.05, \*\**P* < 0.01 by Student’s *t*-test (**C** to **N**) and two-way repeated-measures ANOVA followed by the Sidak test (**C**, **D**, **G** and **H**).

After the mice achieved the criterion in the visual discrimination task, the reward contingency of stimulus images was reversed to investigate the capacity for updating learning based on behavioral flexibility (Reversal learning task; Fig. 6A). The correct response rate started at ∼20% as expected, and gradually increased with repeated training (Fig. 6, G and H). We separated the reversal performance phases into three phases (*23*): the low accuracy phase in the early reversal sessions (early: 0–40% correct response), the chance-level phase around the midpoint of reversal sessions (mid: 40–60% correct), and the high accuracy phase in the late reversal sessions (late: 60–80% correct). *Pcdh17 cKO* mice exhibited a transient performance deficit during the early phase (sessions 3 and 4) (Fig. 6G; main effect, *P* = 0.2346; interaction, *P* = 0.0044) relative to the c*ontrol* mice; however, *Pcdh17 cKO* mice caught up later, and as a result, the total trials and total errors to meet the final criterion are similar between the cKO and control (Fig. 6I).

Consistently, *Pcdh17* cKO mice exhibited increased errors in the early phase but not in the mid or late phases (Fig. 6K). In contrast, *Pcdh10* cKO mice performed similarly to control mice in terms of the percentage of correct responses (Fig. 6H) and total trials and errors to reach the criterion (Fig. 6J). *Pcdh10* cKO mice did not differ in errors in any reversal performance phases compared to control mice (Fig. 6L). These results indicate a slower learning rate in the early phase of reversal learning in *Pcdh17 cKO* mice, but not in *Pcdh10 cKO* mice. We then analyzed perseverative errors (4 consecutive errors were counted as 1 perseverative error) during the early sessions of reversal learning (sessions 1–5), which represent a tendency for persistent and inflexible behavior following an incorrect response (*24*). While *Pcdh10* cKO mice do not show a difference in perseverative errors compared to control mice (Fig. 6N), *Pcdh17* cKO mice made more perseverative errors in sessions 2 and 3 than control mice (Fig. 6M), suggesting that *Pcdh17* cKO mice exhibit reduced behavioral flexibility. Neither *Pcdh17* cKO nor *Pcdh10* cKO mice showed a difference in correct response latency compared to c*ontrol* mice (fig. S12, I and J), indicating that both *Pcdh* cKO mice showed a normal reaction speed to the correct stimulus. Altogether, iSPN-*Pcdh17* cKO mice show impaired task learning and behavioral inflexibility, while iSPN-*Pcdh10* cKO mice show normal learning and flexibility.

### Reduced locomotor and sensory habituation in *Pcdh10* cKO mice, but not in *Pcdh17* cKO mice

We next assessed motor and sensory functions. To examine spontaneous locomotor activity in a novel environment, we performed the open field test. We monitored the distance traveled in the open field box (Fig. 7, A and B). Because the travel distance decreases with habituation to a new environment, the distance traveled was also quantified for the first 5 min (before habituation) and the next 5–15 min (after habituation). All these parameters were similar between *Pcdh17 cKO* mice and *control* mice (Fig. 7A), suggesting that spontaneous locomotor activity and habituation are normal in *Pcdh17 cKO* mice. *Pcdh10 cKO* mice also showed a similar distance traveled for the first 5 min compared to *control* mice. However, *Pcdh10 cKO* mice showed a significantly higher distance traveled during the 5–15 min period (Fig. 7B) than control, suggesting that locomotor habituation in the novel environment was reduced in *Pcdh10 cKO* mice.

**Fig. 7.**
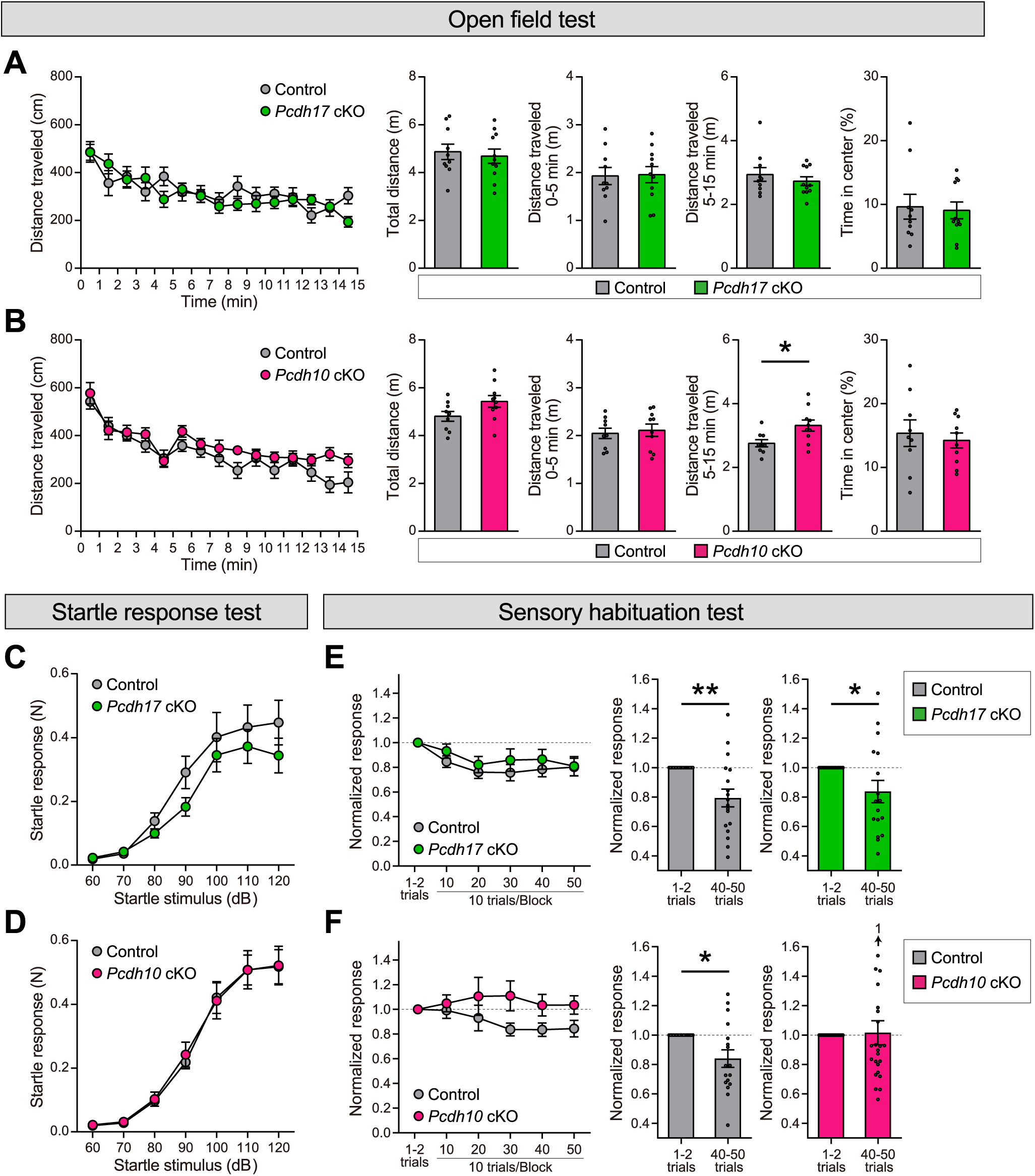
*Pcdh10* cKO but not *Pcdh17* cKO mice show impaired locomotor and sensory habituation. (**A** and **B**) Open field test. The distance traveled in each minute during the 15-min test session, total distance for 15 min, total distance for 0–5 min (before habituation), total distance for 5–15 min (after habituation), and time spent in the center area are shown. The total distance for 5–15 min is significantly increased in *Pcdh10 cKO* mice relative to *control* mice. *n* = 10 mice for *Pcdh17 control*; 11 for *Pcdh17 cKO;* 9 for *Pcdh10 control*; and 10 for *Pcdh10 cKO*. (**C** and **D**) Startle response test. Startle responses for different startle stimuli are similar between *Pcdh17 cKO* and control mice (**C**) as well as between *Pcdh10 cKO* and control mice (**D**). *n* = 11 mice for *Pcdh17 control*; 9 for *Pcdh17 cKO;* 11 for *Pcdh10 control*; and 11 for *Pcdh10 cKO*. (**E** and **F**) Sensory habituation test. Time course of normalized startle responses (left) and the comparison of startle responses at the beginning (1–2 trials) and the end (40–50 trials) of trials (right). *Pcdh17 control*, *Pcdh17 cKO*, and *Pcdh10 control*, but not *Pcdh10 cKO* mice, show decreased normalized responses at the end of trials. *n* = 18 mice for *Pcdh17 control*; 17 for *Pcdh17 cKO;* 17 for *Pcdh10 control*; and 24 for *Pcdh10 cKO*. Data are mean ± SEM. \**P* < 0.05, \*\**P* < 0.01 by Student’s *t*-test.

We then examined sensory habituation of the acoustic startle response. Acoustic startle responses to different startle stimuli were similar between *Pcdh cKO* and *control* mice (Fig. 7, C and D), suggesting that both *Pcdh17 cKO* and *Pcdh10 cKO* mice have a normal response to sensory stimulation. Repeated acoustic stimulation causes a decreased startle response, referred to as “habituation of the acoustic startle response” (*25*, *26*). We found that the startle responses after 40–50 trials in *Pcdh17 control*, *Pcdh17 cKO*, and *Pcdh10 control* mice were significantly lower than those of the initial (the first and second) trials (Fig. 7, E and F), indicating that these mice show sensory habituation of the acoustic startle response. In contrast, *Pcdh10 cKO* mice exhibited a similar level of startle response even after 40–50 trials (Fig. 7F), indicating that they show impaired sensory habituation.

Finally, we investigated whether impaired habituation in *Pcdh10 cKO* mice is due to changes in their anxiety level. In the open field test, time spent in the center area was similar between *Pcdh10* cKO and control mice, as well as between *Pcdh17* cKO and control mice (Fig. 7, A and B; right). In the elevated plus maze, relative to control mice, *Pcdh10* cKO and *Pcdh17* cKO mice spent a similar amount of time in the open arms (fig. S13, A and B). In the light-dark transition test, the time spent in the light area and the transition number were not significantly different between *Pcdh10* cKO and control mice, as well as between *Pcdh17* cKO and control mice (fig. S13, C and D). These results suggest that the anxiety level of *Pcdh10 cKO* mice is normal and that the changes in locomotor and sensory habituation in *Pcdh10 cKO* mice are not due to changes in their anxiety levels. Additionally, prepulse inhibition, in which a weak stimulus given before the startle stimulus suppresses the startle response, was normal in *Pcdh10* cKO and *Pcdh17* cKO mice (fig. S13, E and F), suggesting that sensorimotor gating abilities are normal in both *Pcdh10 cKO* and *Pcdh17 cKO* mice. Taken together, these behavioral experiments revealed that *Pcdh10 cKO* mice, but not *Pcdh17 cKO* mice, exhibit reduced locomotor and sensory habituation.

## Discussion

Here we showed that in the parallel BG circuits, there are specific molecular organizers that establish individual circuits regulating distinct behaviors. We identified PCDH17 and PCDH10 as molecular organizers for parallel indirect BG circuits. In addition to the indirect BG circuits, these PCDHs show complementary expressions in the direct BG circuits (Fig. 1, A and B) as well as in the cortex and thalamus (*10*), suggesting that these PCDHs may organize the entire parallel cortico-BG-thalamo-cortical loop.

### PCDH17 and PCDH10 are molecular organizers for functionally distinct and anatomically segregated parallel BG circuits

Functionally distinct and anatomically segregated parallel neural circuits are thought to underlie parallel information processing in the brain (*27*, *28*). However, how individual circuits are organized was largely unknown, and the molecular mechanisms that establish parallel circuits remain to be elucidated. In the hippocampal circuits, complementary expression of teneurin-3 (Ten3) and latrophilin-2 (Lphn2) has been shown to regulate anatomically segregated circuits (*8*, *29*, *30*) via reciprocal repulsions between these two molecules. However, it is not yet clear whether Ten3 and Lphn2 specify functionally distinct circuits regulating different behaviors. In the BG, it has recently been demonstrated that anatomically segregated BG circuits allow parallel behavioral modulation in mice (*3*). However, how such circuits are established was not known.

In this paper, we showed that PCDH17 and PCDH10 not only anatomically define parallel indirect BG circuits, but also contribute to the functional circuit organization associated with task learning and sensorimotor habituation, respectively. Based on the features of complementary expression and selective homophilic interaction of PCDH17 and PCDH10, parallel BG circuits allow for specific synaptic control, thereby appropriately regulating different behaviors. Thus, PCDH17 and PCDH10 are molecular organizers by which functionally distinct and anatomically segregated parallel circuits are organized for parallel information processing in the brain.

Interestingly, global *Pcdh17* KO mice have been reported to show increased synaptic vesicles in the anterior striatum and inner GPe (*10*), while our study with iSPN-*Pcdh17* cKO mice showed impaired development of synapses. This difference may arise from differences between global and conditional KO models. In global *Pcdh17* KO mice, PCDH17 is deleted in all cell types, potentially triggering compensatory mechanisms that restore or even overcompensate for synaptic phenotypes. In contrast, deleting PCDH17 in specific neurons *(Pcdh17* cKO) may prevent such compensatory responses. In iSPN-*Pcdh17* cKO mice, PCDH17 is absent from the postsynaptic side in the striatum and the presynaptic side in the globus pallidus, creating a mismatch where PCDH17-expressing cells interact with non-expressing neurons. Studies on PCDH19 and other δ2-PCDHs have shown that such mismatched interactions significantly affect cell adhesion and synaptic development (*31–33*). Therefore, in iSPN-specific *Pcdh17* cKO mice, this mismatch may have resulted in a more severe effect than complete deletion.

### Fine-tuning of the parallel indirect BG circuits by PCDHs

PCDH17 and PCDH10 organize parallel indirect BG circuits along the anteroposterior axis in the striatum to the inner-outer axis in the GPe. However, the indirect BG circuits appear more complex – it has been reported that there are 36 domains in the GPe that receive different projections from the striatum (*6*). Thus, additional molecules may exist to fine-tune the parallel indirect BG circuits. Expression analysis of Cadherin superfamily members, including δ1- and δ2-PCDH members, revealed region-specific expression patterns along different axes in the BG (*34*). Their expression patterns are relatively complementary, but there are some overlaps. We also identified a small population of neurons co-expressing *Pcdh17* and *Pcdh10* within BG subregions. This dually-expressing population may have functional implications. First, *Pcdh17*/*Pcdh10* double-positive striatal neurons may preferentially connect with target neurons that also co-express both PCDHs, thereby forming molecularly matched subcircuits. Second, these double-positive neurons might act as bridging populations that interact with both *Pcdh17*-positive and *Pcdh10*-positive neurons, mediating crosstalk between the PCDH17 and PCDH10 pathways. Previous studies have shown that the adhesive properties of δ-PCDHs are tuned by the combinatorial diversity of δ-PCDH expression in neurons (*33*, *35*). Thus, it is possible that the combinatorial diversity of δ-PCDH expressions defines a more detailed parallel circuit structure and produces more diverse circuit functions in the BG.

Additionally, clustered PCDHs encoded by the *Pcdhα*, *Pcdhβ*, and *Pcdhγ* gene clusters are capable of imparting diverse molecular identities at the cell surface through combinatorial expression (*36*). Indeed, it has been shown that patterned clustered PCDH expression in cortical neurons regulates the fine organization of neural circuits (*37*). Clustered PCDHs also regulate dendritic self-avoidance through homophilic repulsion (*38*). In contrast, our results indicate that the loss of PCDH17 and PCDH10 does not disrupt dendritic self-avoidance (i.e., no change in dendritic crossing, fig. S8), suggesting that the function of δ2-PCDHs *in vivo* is not repulsive and differs from that of clustered PCDHs. This highlights the functional diversity of the PCDH family and suggests that multiple PCDH family members may cooperate to fine-tune the organization of parallel BG circuits.

### Potential roles of PCDH17 and PCDH10 in distinct neuropsychiatric disorders

iSPN-*Pcdh17* cKO mice showed deficits in task learning and behavioral flexibility, and iSPN-*Pcdh10* cKO mice showed defects in locomotor and sensory habituation. Are they related to human disorders? In humans, PCDH17 and PCDH10 are implicated in different disorders. The human *PCDH17* gene is associated with mood disorders, including bipolar disorder (BPD) and major depressive disorder (MDD) (*39*). Patients with MDD tend to make poor decisions, which often has a negative impact on their lives (*40*). BPD is associated with impaired decision-making in terms of impulsivity and risk-taking (*41*). In contrast, the human *PCDH10* gene is linked to autism spectrum disorder (ASD) and obsessive-compulsive disorder (OCD) (*42–44*). While the core symptoms of ASD include decreased social motivation, language problems, and restricted behaviors (*45*), many ASD patients also exhibit abnormal sensory sensitivity, described as both hyper- and hypo-reactivity (*46*). Patients with OCD also often exhibit abnormal sensory processing and a reduced ability to filter stimuli that most people do not find bothersome (*47*). While further research is needed, these disease symptoms are consistent with the idea that PCDH17 and PCDH10 mutations may contribute to symptoms of BPD/MDD and ASD/OCD, respectively, through the impairment of distinct indirect BG circuits.

### Limitations of the study and future directions

In this study, we showed that there is no abnormality in the axonal projection patterns of *Pcdh17-* and *Pcdh10-*expressing iSPNs or in the dendritic morphology of iSPNs and dSPNs in *Pcdh17* cKO and *Pcdh10* cKO mice. Therefore, we tentatively conclude that the main defect contributing to the behavioral changes is the synaptic impairment by *Pcdh17-* and *Pcdh10-*expressing iSPNs. However, the abnormal aggregation of SPNs observed in the cKO mice could have secondary effects on circuit function. In addition, there could be other changes that might affect behavioral changes, such as differences in dopamine secretion patterns. Furthermore, A2A-Cre mice can induce recombination not only in the striatum but also in the cortex and hippocampus, which may also contribute to the phenotypes. In future studies, we plan to use optogenetic and chemogenetic strategies to more directly implicate the circuits organized by PCDH17 or PCDH10 in specific behaviors.

## Materials and Methods

### Mice

*Pcdh17* and *Pcdh10* flox-EGFP, cKO EGFP, EGFP knock-in, flox, and cKO mice were generated in our laboratory (see fig. S3 and below). *ACTB-FLPe* (The Jackson Laboratory; JAX:003800) (*48*), *A2A-Cre* (Mutant Mouse Regional Resource Centers; MMRRC_036158) (*11*), *Rosa-STOP-tdTomato* (*49*), *Rbp4-Cre* (Mutant Mouse Regional Resource Centers; MMRRC_037128) (*50*), and *Slc17a7-Cre* (The Jackson Laboratory; JAX:037512) (*50*) mice were described previously. Mice were maintained in standard housing conditions on a 12-hour light/dark cycle with food and water provided *ad libitum*. All animal care and use were in accordance with the institutional guidelines and approved by the Institutional Animal Care and Use Committees at Boston Children’s Hospital and Waseda University.

### Antibodies

Rabbit and guinea pig polyclonal anti-PCDH17 antibodies and rabbit and rat polyclonal anti-PCDH10 antibodies were described previously (*10*, *16*). The PCDH17 and PCDH10 antibodies were used at 1–4 µg/ml. The following primary antibodies were also used: chicken anti-GFP (1:2000; Aves Labs, GFP-1010), mouse anti-NeuN (1:1000; Sigma-Aldrich; MAB377), guinea pig anti-VGAT (1:1000; Nittobo Medical; VGAT-GP-Af1000), guinea pig anti-D1R (1:1000; Nittobo Medical; D1R-GP-Af500), rat anti-D1R (1:1000; Sigma-Aldrich; D2944), rabbit anti-D2R (1:1000; Nittobo Medical; D2R-Rb-Af960), rabbit anti-dsRed (1:500; Clontech; 632496), guinea pig anti-VGLUT1 (1:5000; EMD Millipore; AB5905), guinea pig anti-VGLUT2 (1:1000; Nittobo Medical; VGluT2-GP-Af810), rabbit anti-VGAT (1:1000; Synaptic Systems; 131003), mouse anti-PSD95 (1:1000; NeuroMab; 75-028), and mouse anti-Gephyrin (1:500; Synaptic Systems; 147021). The following secondary antibodies (Thermo Fisher Scientific) were used for immunohistochemistry: Donkey anti-Rabbit IgG, Alexa Fluor 488 (A-21206), Goat anti-Rabbit IgG, Alexa Fluor 488 (A-11034), Alexa Fluor 568 (A-11036), Donkey anti-Mouse IgG, Alexa Fluor 488 (A-21202), Goat anti-Guinea Pig IgG, Alexa Fluor 488 (A-11073), Alexa Fluor 568 (A-11075), Alexa Fluor 647 (A-21450), Goat anti-Chick IgY, Alexa Fluor 488 (A-11039), Donkey anti-Rat IgG, Alexa Fluor 488 (A-21208), Alexa Fluor 555 (A-78945). Additionally, Goat anti-Mouse IgG, Alexa Fluor 647 (Jackson ImmunoResearch; 115-605-146) was used.

### Generation of *Pcdh17* and *Pcdh10* conditional knock-out mice

*Pcdh17* and *Pcdh10 flox-*EGFP mice were generated using the CRISPR/Cas9 system. Two gRNAs were designed to remove the 1st exon of *Pcdh17* or *Pcdh10* from the WT allele (see fig. S3). The target sequences of *Pcdh17* gRNAs were 5’-AGGTTCAAACTTGCGTGATG*AGG*-3’ and 5’-TGACCCTGTGTGAATACCAT*TGG*-3’ (the PAM sequence is in italic). The target sequences of *Pcdh10* gRNAs were 5’-CTCCTTTATTCCGACAGTGT*TGG*-3’ and 5’-GAGGGGCCAATCACTGACAA*AGG*-3’. Sigma-Aldrich provided the crRNA and tracrRNA. The donor vector, with a pSV backbone (*51*), contained the loxP-flanked 1st exon of the *Pcdh* gene and the FRT-flanked EGFP-polyA cassette, with 5’ and 3’ homology arms spanning ∼4 kb for homology-directed repair. The pSV backbone vector was described previously (*51*). The donor vector, gRNAs, crRNA, and tracrRNA were injected with Cas9 protein (PNA Bio) into the pronucleus of fertilized zygotes (C57BL/6N), and the injected zygotes were transferred into the oviducts of pseudo-pregnant female mice. The positive F0 pups having a flox-EGFP allele were identified by the detection of the 5’ arm and 3’ arm of the *Pcdh17* and *Pcdh10* genes by genomic PCR, respectively. *Pcdh* flox-EGFP mice were crossed with *ACTB-FLPe* mice to remove the EGFP-polyA cassette through FRT/FLP-mediated excision to generate *Pcdh* flox mice. To generate constitutive *Pcdh-EGFP* knock-in mice via germline recombination, we used either *Rbp4-Cre* or *Slc17a7-Cre* mice to remove the first exon of each *Pcdh* allele. These Cre lines have been reported to cause germline recombination when inherited from the father⁵⁰. Germline recombination was confirmed in offspring from *Rbp4-Cre/Pcdh17^floxE/+^* and *Slc17a7-Cre/Pcdh10^floxE/+^*male mice (see fig. S3, E and F), resulting in the generation of *Pcdh17^EGFP/+^*and *Pcdh10*^EGFP/+^ mice. *Pcdh17* wild-type and conditional alleles (flox-EGFP and flox) were identified using PCR assays in which primers P17Fw (5’-CCTAGCGGGAGTGAGGAAAACCAAGC-3’) and P17Rv (5’- CAAAGCCTTTAGCTTGCCTTCAGCCG-3’) amplified a wild-type fragment and a 5’ loxP-inserted fragment, respectively (see fig. S3C). *Pcdh10* wild-type and conditional alleles were detected using PCR assays in which primers P10Fw (5’-CCCATCAGCAGGAAGACAAACCTAGG-3’) and P10Rv (5’-

AGAGCGTCTCCAAATCGAGCCTCATT-3’) amplified a wild-type fragment and a 5’ loxP-inserted fragment (see fig. S3D). The presence of the EGFP sequence was confirmed by PCR using primers GFP-Fw (5’-ATCTTCTTCAAGGACGACGGCAACTACAAG-3’) and GFP-Rv (5’-AGAGTGATCCCGGCGGCGGTCACGA-3’), which yielded a single band. *Pcdh17^EGFP/+^* and *Pcdh10*^EGFP/+^ mice were identified by the presence of both the EGFP sequence and the wild-type *Pcdh* allele, and the absence of the 5′ loxP-inserted fragment, as determined by PCR using the primers described above (see fig. S3, E and F). *Pcdh* cKO mice were generated by mating *Pcdh* flox mice with *A2A-Cre* mice. Immunostaining confirmed decreases in PCDH17 and PCDH10 proteins in *Pcdh* cKO mice (see fig. S5, A and B). *Pcdh17* and *Pcdh10* flox-EGFP and flox mice were backcrossed with C57BL/6J mice at least 5 times and maintained on a C57BL/6J background. To minimize experimental variability, we used littermates as controls. Specifically, we crossed *Pcdh^flox/+^* males with *Pcdh^flox/+^* females to obtain *Pcdh^flox/flox^* and *Pcdh^+/+^* mice within the same litter for comparison.

### AAV preparation and retro-orbital injections for sparse labeling of SPNs

pAAV-hSyn-EGFP-CAAX (Addgene; Plasmid #208684) was used to produce recombinant AAVs expressing membrane-tagged EGFP (mEGFP). AAVs of serotype PHP.eB were produced. AAVpro 293T cells (Takara Bio) were transfected with pAAV-hSyn-EGFP-CAAX, pAdDeltaF6 (Addgene; Plasmid #112867), and pUCmini-iCAP-PHP.eB (Addgene; Plasmid #103005) using polyethylenimine. 5 days after transfection, the cells and media containing AAVs were collected, and the cells were lysed. After removing cell debris, AAVs were purified by ultracentrifugation and suspended in PBS. AAV titers were determined by real-time quantitative PCR. The titer of the viruses used for injection in this study was 1–5 × 10¹² genome copies (GC)/ml.

For AAV retro-orbital injections, 6- to 9-week-old mice were anesthetized with a mixture of three anesthetics, including medetomidine hydrochloride (0.3 mg/kg), midazolam (4 mg/kg), and butorphanol tartrate (5 mg/kg). A total of 100 μl of AAV suspended in PBS was injected into the retro-orbital sinus using a 30G needle. Mice were analyzed 2–3 weeks after AAV infection.

### Immunohistochemistry

Male and female mouse brains were perfused with 4% paraformaldehyde (PFA) in PBS, post-fixed in 4% PFA in PBS overnight at 4°C, and then cryoprotected in 30% sucrose in PBS at 4°C. For PCDH17 and PCDH10 staining, mouse brains were perfused with 2% PFA in PBS without post-fixation. Brains were frozen and sectioned at 16 μm thickness on a CM3050S or CM1950 cryostat (Leica). For PSD95 staining, sections were pretreated with 1 mg/ml pepsin in 0.2 N HCl for 5 min at room temperature before staining. To enhance the dendrite signal for synaptic puncta analysis, sections were co-stained with anti-dsRed antibodies. Sections were incubated with blocking buffer (2% BSA/2% goat serum/0.1% Triton X-100 in PBS) for 1 hour at room temperature, incubated with primary antibodies in blocking buffer overnight at 4°C, and incubated with fluorescent secondary antibodies (1:500; Thermo Fisher Scientific) in blocking buffer for 2 hours at room temperature. Sections were washed with PBS between incubation steps. Sections were mounted with Fluoromount-G.

### Imaging

#### Imaging of PCDH17 and PCDH10 staining

Images were taken with a BZ-X810 fluorescence microscope (Keyence), an LSM700 confocal microscope (Zeiss), or an FV3000 confocal microscope (Evident). Images were obtained with the identical acquisition settings regarding the exposure time, laser power, detector gain, and amplifier offset. For wide-field imaging, images were obtained by stitching several adjacent images together. The average signal intensity was quantified using ImageJ. The ROIs analyzed in fig. S5 are: the total area of the anterior and posterior striatum, inner and outer GPe, and posterior SNr (PCDH17; fig. S5, A and C), and the total area of the anterior and posterior striatum, inner and outer GPe, and anterior SNr (PCDH10; fig. S5, B and D). The signal intensities in the corpus callosum for the striatum, internal capsule for the GPe, and cerebral peduncle for the SNr were calculated as the background signals. The background signals were subtracted from the signal intensities in the ROIs. For fig. S6, A and B, single-plane high-resolution images for PCDH17/PCDH10, D1R, and D2R in the anterior and posterior striatum were acquired at a resolution of 1024 × 1024 pixels using a 100× objective lens with 1× zoom. The cell bodies (ROIs) of D1R-positive dSPNs and D2R-positive iSPNs were identified based on the D1R/D2R signal distributions¹³. The average intensities of PCDH17 or PCDH10 in the anterior and posterior striatum were quantified within the ROIs. Signal intensities in the nuclei of these cells were considered as background and were subtracted from the signal intensities within the ROIs.

#### Quantification of Pcdh17 and Pcdh10 mRNA expression

Double fluorescent *in situ* hybridization images were analyzed using ImageJ. The signal intensities in the corpus callosum for the quantification in the striatum, internal capsule for GPe/GPi, and cerebral peduncle for SNr were calculated as the background signals. Background-subtracted binary images were prepared. To assess co-expression, *Pcdh17* and *Pcdh10* binary images were overlaid. Cells were manually counted as double-positive if the overlap between *Pcdh17* and *Pcdh10* signals occupied more than 10% of the total signal area. Otherwise, *Pcdh*-positive cells were categorized as either *Pcdh10*-only or *Pcdh17*-only. The percentages of each cell type were calculated, and the results were visualized as pie charts. The numbers of cells analyzed were: striatum (1168 cells from 2 fields), GPe (709 cells from 3 fields), GPi (187 cells from 2 fields), and SNr (234 cells from 1 field).

#### Imaging PCDH17^+^ and PCDH10⁺ inhibitory terminals in the GPe

Sagittal brain sections from *Pcdh17^EGFP/+^* and *Pcdh10^EGFP/+^*mice were immunostained with antibodies against EGFP, VGAT, and NeuN to visualize inhibitory terminals and GPe neurons. Images were taken with an FV3000 confocal microscope (Evident). Images were obtained with the identical acquisition settings regarding the exposure time, laser power, detector gain, and amplifier offset. For fig. S4, A and C, single-plane images were acquired at a resolution of 1024 × 1024 pixels using a 20× objective lens with 1× zoom; for fig. S4, B and D, images at 1024 x 1024 pixels were acquired as a z-stack (8 optical sections, 0.43 μm step size) using a 60x objective lens with a 2× zoom, and subjected to maximum intensity projection. Synaptic targeting in the GPe was quantified with ImageJ. Background intensity for VGAT and NeuN was determined using the Auto Threshold plugin. For EGFP, signal intensities within the nuclei of PCDH-negative cells were used to define background levels. Background-subtracted binary images were prepared for all channels. VGAT binary images were overlaid onto EGFP binary images, and EGFP⁺/VGAT⁺ puncta, representing PCDH-positive inhibitory terminals, were extracted. NeuN signals were also binarized to define the neuronal cell body. The number of EGFP⁺/VGAT⁺ puncta per EGFP⁺/NeuN⁺ and EGFP^-^/NeuN^+^ GPe cell body was quantified using ImageJ’s “Analyze Particles” function. Puncta smaller than 5 pixels were excluded from analysis.

#### Imaging of D1R and D2R staining

Images were taken with an LSM700 confocal microscope (Zeiss) or an FV3000 confocal microscope (Evident). For each antibody, images were obtained with the identical acquisition settings regarding the exposure time, laser power, detector gain, and amplifier offset. For Fig. 2, A to D, single-plane images were acquired at a resolution of 1024 × 1024 pixels using a 25× objective lens with 1× zoom; for fig. S7, E to H, single-plane images were acquired at a resolution of 1024 × 1024 pixels using a 10× objective lens with 1× zoom; and ROIs corresponding to the anterior and posterior striatum were subsequently defined. For each image, the green (D2R) and red (D1R) channels were separated. For each channel, the background was corrected by subtracting the average intensity of three random points from the entire image using ImageJ. The random points were chosen using an online random number generator, which provided three pairs of random numbers between 1 and 1024, where each pair was used as the x and y coordinates. If a point fell into an area with no signal, it was replaced with another random point. The pixel intensity arrays were then exported to finish the analysis using Python. Specifically, for the D1R array, the data was smoothed by averaging the intensities over squares measuring 5 × 5 pixels. The averaged intensities were then normalized by setting the maximum intensity to 1, the minimum intensity to 0, and scaling all the other intensities accordingly. The array was then flattened, with each row being concatenated with the previous and following rows, creating a 1-D vector containing all the rows of the D1R array. These steps were then repeated with the D2R array. The similarity coefficient was calculated by taking the normalized dot product of the two vectors, which measures how correlated the D1R and D2R intensities are in each 5 × 5 region. The contour maps were plotted in Python using matplotlib.

For fig. S7, A to D, single-plane images were acquired at a resolution of 1024 × 1024 pixels using a 60× objective lens with 1× zoom. To measure the nearest neighbor distance, dSPNs and iSPNs were visually identified based on D1R and D2R signal distribution around the cell bodies. The cell bodies were then manually plotted using Illustrator (Adobe Systems). Nearest neighbor distances were quantified using the BioVoxxel Toolbox plugin in ImageJ, and the average nearest neighbor distance per image was calculated.

#### Imaging of cell morphologies

For the imaging of cell morphologies of dSPNs and iSPNs in the striatum, mice that underwent retro-orbital injections were perfused with 4% PFA in PBS. Brains were removed, post-fixed in 4% PFA in PBS overnight at 4°C, and cryoprotected in 30% sucrose in PBS at 4°C. 100 μm-thick sections were cut on a CM1950 cryostat and collected in PBS. Sections were washed with PBS and then mounted with Fluoromount-G.

For imaging of cell bodies and dendrites of dSPNs and iSPNs, images of mEGFP and tdTomato fluorescence were taken with an FV3000 confocal microscope from the anterior and posterior striatum. Images at 512 × 512 pixels were acquired as a z-stack (30 optical sections, 1.5 μm step size) using a 20× objective lens with a 1× zoom. Z-stack images were subjected to maximum projection. Images were obtained with the identical acquisition settings regarding the exposure time, laser power, detector gain, and amplifier offset. dSPNs and iSPNs were determined by whether the cell body signal of mEGFP in single-plane images was tdTomato^+^ (iSPNs) or tdTomato^-^ (dSPNs). The cell body size (area) of dSPNs and iSPNs was measured using ImageJ. For the analysis of dendritic crossings, maximum intensity images were inverted and contrast-enhanced using Photoshop (Adobe Systems). Dendritic crossings in single SPNs were manually quantified by counting the number of branch overlaps.

#### Imaging of axon terminals

In *A2A-Cre;Rosa^STOP-tdT^;Pcdh^flox-EGFP^* mice, the EGFP signal inside the GPe visualizes the area of axonal innervation from *Pcdh*-expressing iSPNs. Sagittal sections were stained for EGFP, and whole brain images were obtained by stitching several adjacent images together using a BZ-X810 fluorescence microscope (Keyence) with 4x lens. Image analysis was performed using ImageJ. The signal intensity of the internal capsule was calculated as the background signal and subtracted from that of the GPe. Analysis of axon terminals was done using a 50-pixel wide line scan crossing the GPe (see fig. S9). To consistently localize this line scan, the following steps were taken: The DAPI signal was used to find the granule cell layer of the dentate gyrus (DG). Hereafter, the apex and the anterior end of the upper blade of the DG were connected with a straight line (fig. S9, B and C). Next, a parallel line to the first one was drawn through the geometrical center of the GPe, which was determined using the tdTomato signal (fig. S9, A to D). This line was then used to perform a line scan oriented from the anterior side of the GPe (closer to the striatum) to the posterior side of the GPe (fig. S9, E and F). The highest axonal density was set as 1 and the remaining values were normalized accordingly. Line scans were performed on 2–4 images per mouse and then averaged. Thereafter, the EGFP signal represented by the average line scan was evaluated by introducing a threshold intensity value.

For each mouse, the average EGFP intensity of the medial geniculate nucleus (MGN) was multiplied by the factor 1.5 (fig. S9E, white box), normalized according to the values obtained in the line scan, and used as an individual threshold value defining the border of axonal innervation. At this intensity value, on the y-axis of the line scan, a horizontal bar was introduced. Doing so yielded two crossing points with the average EGFP-line scan (fig. S9G). The distance from the beginning of the line scan to the first crossing point (Distance 1), the distance between the two crossing points (Distance 2), and the distance between the second crossing point and the posterior edge of the GPe (end of line scan, Distance 3) were determined and used to compare between the genotypes (Fig. 2, E to H). To evaluate the density of axon terminals in the GPe (fig. S9, H and I), we measured the EGFP intensity in the GPe and divided it by the EGFP intensity in the striatum. For this purpose, in iSPN-*Pcdh17* cKO mice we measured intensity in the anterior part of the striatum and the inner region of the GPe; for iSPN-*Pcdh10* cKO mice, the posterior part of the striatum and the posterior part of the GPe were assessed.

#### Imaging of synaptic puncta

Images were taken with an LSM700 confocal microscope (Zeiss). For each individual antibody, images were obtained with the identical acquisition settings regarding the exposure time, laser power, detector gain, and amplifier offset. In the striatum, images at 1024 × 1024 pixels were acquired as a z-stack (5 optical sections, 0.5 μm step size) using a 63× objective lens with a 1.5× zoom. Background intensity was removed by using ImageJ’s ‘rolling ball’ algorithm. For VGLUT1, VGLUT2, and PSD-95 in the striatum, and VGAT in the GPe, puncta on dendrites or axons were quantified using the ‘Synapse on Dendrite’ quantifier plugin for ImageJ (*52*). Thereby, puncta density (number of puncta/dendrite area) and puncta size were determined. For the analysis of VGAT and Gephyrin puncta in the striatum, the number of synaptic puncta per tdTomato^+^ cell body, along with the puncta size, were calculated using the ‘Synapse on Dendrite’ plugin. In the GPe, z-stack settings were adjusted to 12 optical sections with 0.5 μm step size. Background-subtracted images were prepared, and the density and size of puncta were quantified using ImageJ’s ‘analyze particle’ function. Puncta smaller than 5 pixels were excluded from analysis.

### Electrophysiology

All electrophysiological experiments and analyses were done blind. Recordings were done on male and female mice from ages P12–P18. Mice were decapitated, brains were quickly removed, and then 300 µm parasagittal slices were cut using a VT1200S vibratome (Leica). Slices were cut in an ice-cold cutting solution containing: 206 mM sucrose, 2.8 mM KCl, 2 mM MgSO_4_, 1 mM MgCl_2_, 1.25 mM NaH_2_PO_4_, 1 mM CaCl_2_, 10 mM glucose, 26 mM NaHCO_3_, and 0.4 mM ascorbic acid. After cutting, slices were moved into room temperature artificial cerebral spinal fluid (aCSF) containing: 125 mM NaCl, 2.5 mM KCl, 2 mM CaCl_2_, 1 mM MgCl_2_, 1.25 mM NaH_2_PO_4_, 16.6 mM glucose, 26 mM NaHCO_3_. Recording was then conducted in the same aCSF, at least one hour after cutting. All solutions were continuously bubbled with 95% oxygen/5% carbon dioxide. Neurons were visualized using a customized Scientifica/Olympus microscope.

Data were obtained with a Multiclamp 700B amplifier (Molecular Devices), digitized with a Digidata 1440A (Molecular Devices), and collected with Clampex 10.7 (Molecular Devices). Whole-cell patch-clamp recordings were conducted with 4-6 MΩ pipette containing: 120 mM CsMeSO_4_, 15 mM CsCl_2_, 8 mM NaCl, 0.2 mM EGTA, 2 mM Mg-ATP, 0.3 mM Na-GTP, 10 mM TEA, 10 mM HEPES, 5 mM QX-314 (to block sodium channels), and 0.15% lucifer yellow (to fill the cells). The pH of the internal solution was adjusted to 7.3 with CsOH. For sEPSCs, cells were held at -70 mV. The aCSF was supplemented during recordings with 50 µM picrotoxin (to block inhibition) and warmed to 32°C. Finally, sEPSCs were analyzed using Minianalysis (Synaptosoft).

To record sIPSCs in the GPe, sagittal sections were prepared as above. We used the same aCSF (supplemented with 25 µM DL-APV and 10 µM CNQX to block excitation) and internal solutions. Neurons were targeted along the edge of the GPe (in the putative PCDH10^+^ region) based upon tdTomato fluorescence. Neurons were held at 10 mV, and sIPSCs were recorded.

For short-term plasticity, 300 μm parasagittal sections were cut from P11-P23 mice as described above in an ice-cold solution containing 110 mM choline chloride, 25 mM NaHCO₃, 2.5 mM KCl, 7 mM MgCl₂, 0.5 mM CaCl₂, 1.25 mM NaH_2_PO_4_, 25 mM glucose, 11.6 mM ascorbic acid, and 3.1 mM pyruvic acid. Then slices were transferred to a solution containing 125 mM NaCl, 2.5 mM KCl, 2 mM CaCl₂, 1 mM MgSO₄, 1.25 mM NaH_2_PO_4_, 26 mM NaHCO₃, and 20 mM glucose. During recording, this solution was also supplemented with 50 μM picrotoxin. Cells were patched in the anterior striatum with an internal solution containing 135 mM CsCl, 10 mM HEPES, 1 mM EGTA, 1 mM Li-GTP, 4 mM Mg-ATP, 5 mM MgCl₂. Cells were held at -70 mV. Signals were filtered at 3 kHz and digitized at 20 kHz. Two micropipettes were filled with 3 M NaCl and were placed in the cortex at the border of the striatum. The paired pulse ratio was measured with interstimulus intervals of 20, 30, 50, 70, 100, and 200 ms. To measure short-term depression, EPSCs were evoked by 200 stimuli at 10 Hz.

### Behavioral tests

All data acquisition and analyses were carried out by an individual blind to the genotype. 2- to 4-month-old male mice were used for experiments. Because female mice have estrous cycles that may result in more behavioral variability than male mice (*53*), we used male mice in the behavioral studies.

#### Open field test

Mice were placed in the center of an open-field apparatus (41 × 41 × 38 cm (W × D × H)) illuminated at 30 lux and allowed to move freely for 15 min. Distance traveled for each 1 min, total distance traveled, distances traveled for the first 5 min and the next 10 min, and time spent in the center area (20 x 20 cm; about 25% of the total area) were recorded and analyzed with EthoVision XT software (Noldus).

#### Acoustic startle response, sensory habituation, and prepulse inhibition tests

The startle reflex measurement system (Kinder Scientific) was used for assessing acoustic startle responses, habituation of the acoustic startle response, and prepulse inhibition. A test session began by placing a mouse in a chamber, where it was left undisturbed for 5 min. The startle response was recorded for 250 ms starting with the onset of a startle stimulus. The background noise level in the chamber was 60 dB. The peak startle amplitude during the 250 ms sampling window was recorded as a measure of the startle response. The average intertrial interval was 15 s (range, 10–20 s). The acoustic startle response test was done on day 1. The intensities of startle stimuli were 60, 70, 80, 90, 100, 110, and 120 dB. The habituation test of the acoustic startle response was done on day 2. The 110 dB startle stimuli were applied 50 times. Normalized startle responses were determined by dividing each response by the mean of the first two responses. For the analysis, 10 normalized startle responses were binned. For the quantitative assessment of the habituation, the average of the last 20 startle responses was divided by the average of the first two responses. The prepulse inhibition test was done on day 3. The prepulse sound was presented 100 ms before the startle stimulus (110 dB), and its intensity was 70, 74, 78, or 82 dB. Each trial was presented in pseudorandom order. The following formula was used to calculate the % prepulse inhibition of the startle response: 100 - [100 × (startle response on prepulse trials/startle response on 110 dB startle trials)].

#### Elevated plus maze test

The elevated plus maze consisted of two open arms and two closed arms of the same size (35 × 6 cm) extending from the central area (6 × 6 cm). The maze was elevated 75 cm from the ground and illuminated at 3 lux. Mice were placed in the central square of the maze facing one of the open arms and allowed to move for 10 min. Times spent in the open arm, closed arm, and center area were recorded and analyzed with EthoVision XT software.

#### Light-dark transition test

The light-dark box consisted of two compartments: a white (light) box without a lid (500 lux) and a black (dark) box with a lid (both 25 × 25 × 25 cm). The two boxes were separated by a vertical sliding door that remained open (4 × 6 cm). Mice were placed in the black box and allowed to move between boxes for 10 min. Times spent in the light box and dark box, and the transition numbers between boxes were recorded and analyzed with EthoVision XT software.

#### Behavioral experiments with a touchscreen operant system

The test was conducted in a touchscreen operant chamber equipped with an infrared touchscreen, a black mask with two windows, a liquid dipper, and a reward tray (Lafayette Instrument Company). Mice could access 10% condensed milk (Borden Sugar Condensed Eagle Brand Milk) as a reward from the reward tray. The operant chamber is controlled using ABET II (Lafayette Instrument Company) and Whisker (Lafayette Instrument Company) software.

Throughout the testing, the mice’s body weights were maintained at 85% of free-feeding weight by restriction feeding. During restriction feeding, mice could access around 2 g of regular mouse chow per day.

#### Pre-training

Pre-training involved three stages. Mice were required to meet a set criterion for each stage before proceeding to the next stage. In stage 1 (acclimation), mice were placed in the chamber for 60 min and exposed to 200 µl of condensed milk in the reward tray. Once the mice had consumed all the milk in 60 minutes, they proceeded to the next stage. In stage 2 (reward acquisition training), the mice were trained to collect a milk reward (20 μl each time) from the reward tray after the following cues: a high-pitched tone and illumination of the tray light. After the reward was collected, the tray illumination was turned off, and a 30 sec inter-trial interval was initiated before the delivery of the next reward. Mice that successfully completed 40 trials in 60 min proceeded to the next stage. In stage 3 (stimulus touch training), mice were trained to touch visual stimuli that were randomly presented in one of the two spatial locations on the screen. 40 different shapes were used for the visual stimuli, none of which were used for the visual discrimination or reversal learning task. If mice touch the blank screen, no reward is given. Touching the visual stimulus results in its disappearance, accompanied by the delivery of the reward, a high-pitched tone, and tray illumination. Reward collection turned off the tray illumination and initiated the next trial. Mice that successfully completed 60 trials in 60 min were considered to have met the criterion in stage 3. The days to the criterion were quantified for stages 1–3. Touch latency (time from visual stimulus presentation to touch response) for stage 3 and reward latency (time from reward delivery to reward retrieval) for stages 2 and 3 were measured. The median of reward latency and touch latency for total trials were calculated using the ABET II analysis program, and the average of those values for each mouse group was plotted on the graph.

*Visual discrimination task* (see Fig. 6, A and B). In the visual discrimination task, mice were trained to touch the correct visual stimulus from two different visual stimuli that were randomly presented on the screen. Images of a fan and marbles were used for the visual stimuli. The correct visual stimulus was counterbalanced in half of the mouse groups. Touching the correct visual stimulus (correct response) results in its disappearance, accompanied by the delivery of the reward, a high-pitched tone, and tray illumination. Reward collection turned off the tray illumination and initiated the next trial. Touching the incorrect visual stimulus (incorrect response) results in a 5 sec timeout accompanied by a low-pitched tone and the illumination of a house light. After the timeout, the tray light illuminated so that the mouse could nose-poke the tray to initiate the next trial. One session was conducted per day and lasted 60 min, with a maximum of 60 trials. Mice that successfully achieved an average correct response rate of at least 80% in 60 trials over two consecutive days were considered to have met the criterion in the visual discrimination task. At least four sessions were completed, regardless of whether this criterion was met. Percentage of correct responses (correct responses/60 trials), total trials required to criterion, total errors made to criterion, and correct response latency (time from visual stimulus presentation to correct touch response) were measured. In Fig. 6, C and D (correct response) and fig. S12, G and H (correct response latency), because mice that met the criterion earlier than day 9 were immediately moved to the reversal learning task, the values when the mice met the criterion were repeatedly plotted until day 9 in the graphs. The median of correct response latency was calculated using the ABET II analysis program.

#### Reversal learning task

In the reversal learning task, the order of the task was the same as in the visual discrimination task, but for each mouse, the correct visual stimulus was switched to the other visual stimulus. One session was conducted per day and lasted 60 minutes, with a maximum of 60 trials. However, because a few mice had a large decrease in trial number per day in the early stages of reversal learning, we combined trials from the following day to adjust the number of trials to 60 per session. Mice that successfully achieved an average correct response rate of at least 80% in 60 trials over two consecutive days were considered to have met the criterion for reversal learning. Regardless of whether this criterion was met, at least 13 sessions were completed. In addition to the analysis in the visual discrimination task, total errors in the early (correct response 0–40%), mid (correct response 0–60%), and late (correct response 60%–criterion) phases were determined. Perseverative errors for sessions 1–5 were also measured, where four consecutive incorrect responses were counted as 1 perseverative error.

### Statistics and reproducibility

Statistical analyses were performed using GraphPad Prism software. The statistical tests performed were two-tailed Student’s *t*-test, Mann-Whitney U test, two-way ANOVA followed by the Sidak test, and two-way repeated-measures ANOVA followed by the Sidak test as indicated in the results and figure legends. All data are expressed as mean ± SEM. Sample sizes (*n*) are indicated in the figure legends. Significance was set as \**P* < 0.05, \*\**P* < 0.01, \*\*\**P* < 0.001 or \*\*\*\**P* < 0.0001 for all data. The sample sizes were not predetermined by statistical methods; however, our sample sizes were similar to those reported in previous publications in the field (*10*, *52*, *54*). Steps in the experiments were randomized to minimize the effects of confounding variables, including how mice were chosen for experiments, order of treatments, etc. Imaging was done in the same fashion among conditions. For quantification, cells and fields from brain sections were chosen randomly from the region of interest. No data points were excluded from any experiments.

## Acknowledgments

We thank Sivapratha Nagappan-Chettiar, Akiko Terauchi, Mariel Seiglie, and Masahiro Yasuda for feedback on the manuscript. We thank Yuichi Makino for technical assistance with the touchscreen operant system and Elizabeth Strang for help with behavioral tests. We thank Naoya Noguchi and Chloe DiScipio for mouse colony management, and Ryo Shibata, Takuto Miwa, and Nodoka Hayakawa for AAV preparation. We also thank the Mouse Gene Manipulation Core and the Animal Behavior and Physiology Core at Boston Children’s Hospital, as well as the Department of Statistics at Harvard University.

## Funding

This work is supported by the NIH grant MH125162 (H.U.), SFARI grant 568285 (H.U.), Harvard Brain Science Initiative Bipolar Disorder Seed Grant (H.U.), JSPS KAKENHI Grant Number JP24K02133 (N.H.), the Uehara Memorial Foundation (N.H.), the Brain Science Foundation (N.H.), the Naito Foundation (N.H.), the Astellas Foundation for Research on Metabolic Disorders (N.H.), Takeda Science Foundation (N.H.), and the Boehringer Ingelheim Fonds (J.M.B.).

## Author contributions

Conceptualization, N.H. and H.U.; Methodology, N.H., J.M.B., E.M.J.-V., A.D., and H.U.; Investigation, N.H., J.M.B., E.M.J.-V., M.H., K.M., A.D., V.R.R., J.S., and T.I.; Writing – original draft, N.H. and H.U.; Writing – review and editing, N.H., J.M.B., E.M.J.-V., and H.U.; Funding Acquisition, N.H., H.U.; Supervision, H.U.

## Competing interests

The authors declare no competing interests.

## Data and materials availability

All data and materials supporting the findings of this study are included in the main text and Supplementary Information.

---

**Fig. S1.**
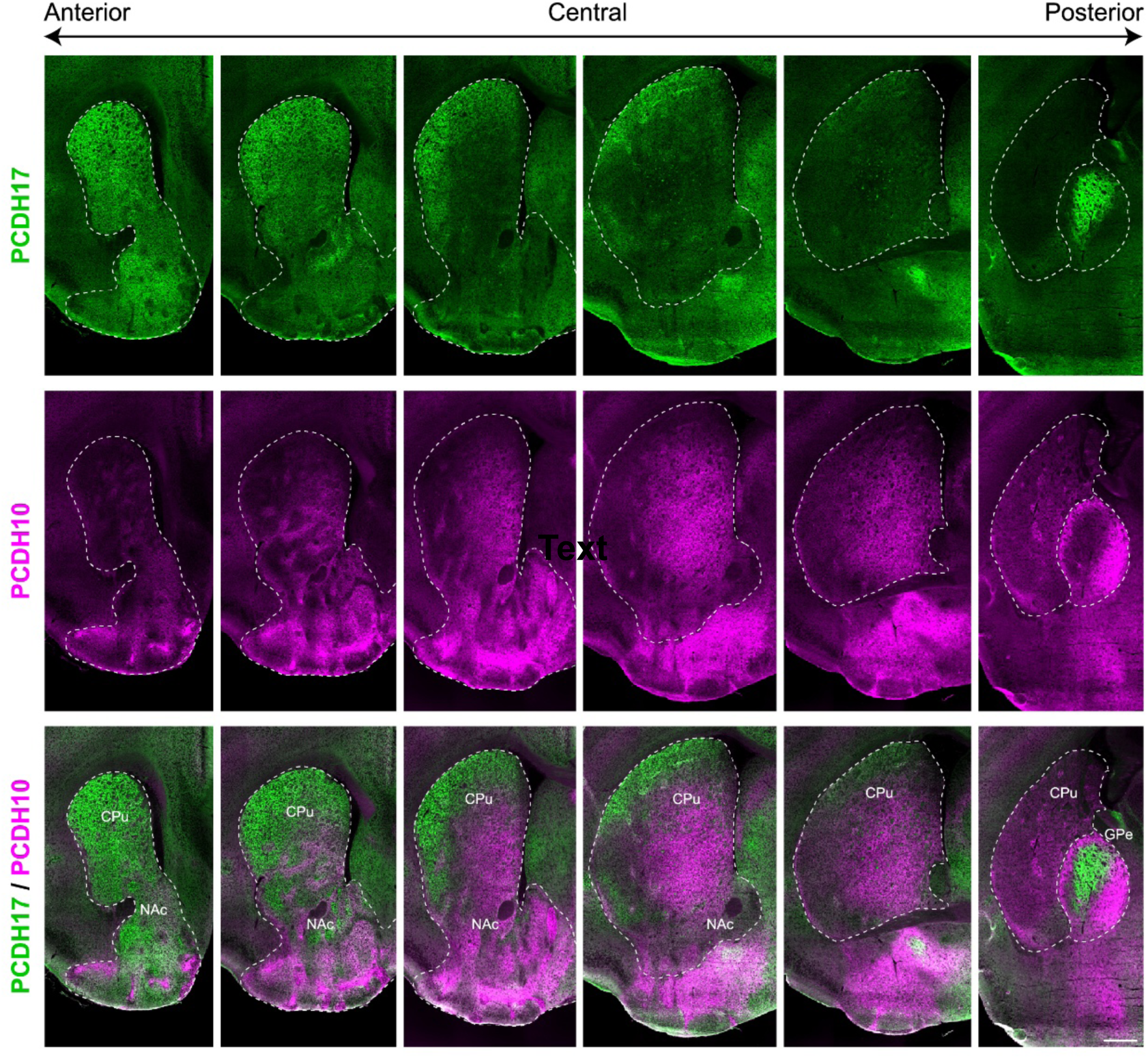
Complementary expression patterns of PCDH17 and PCDH10 proteins in the striatum. Double staining for PCDH17 (green) and PCDH10 (magenta) of coronal striatal sections from P11 mice. PCDH17 and PCDH10 proteins show complementary expression patterns in the striatum, including the caudate putamen (CPu) and nucleus accumbens (NAc). The scale bar represents 0.5 mm.

**Fig. S2.**
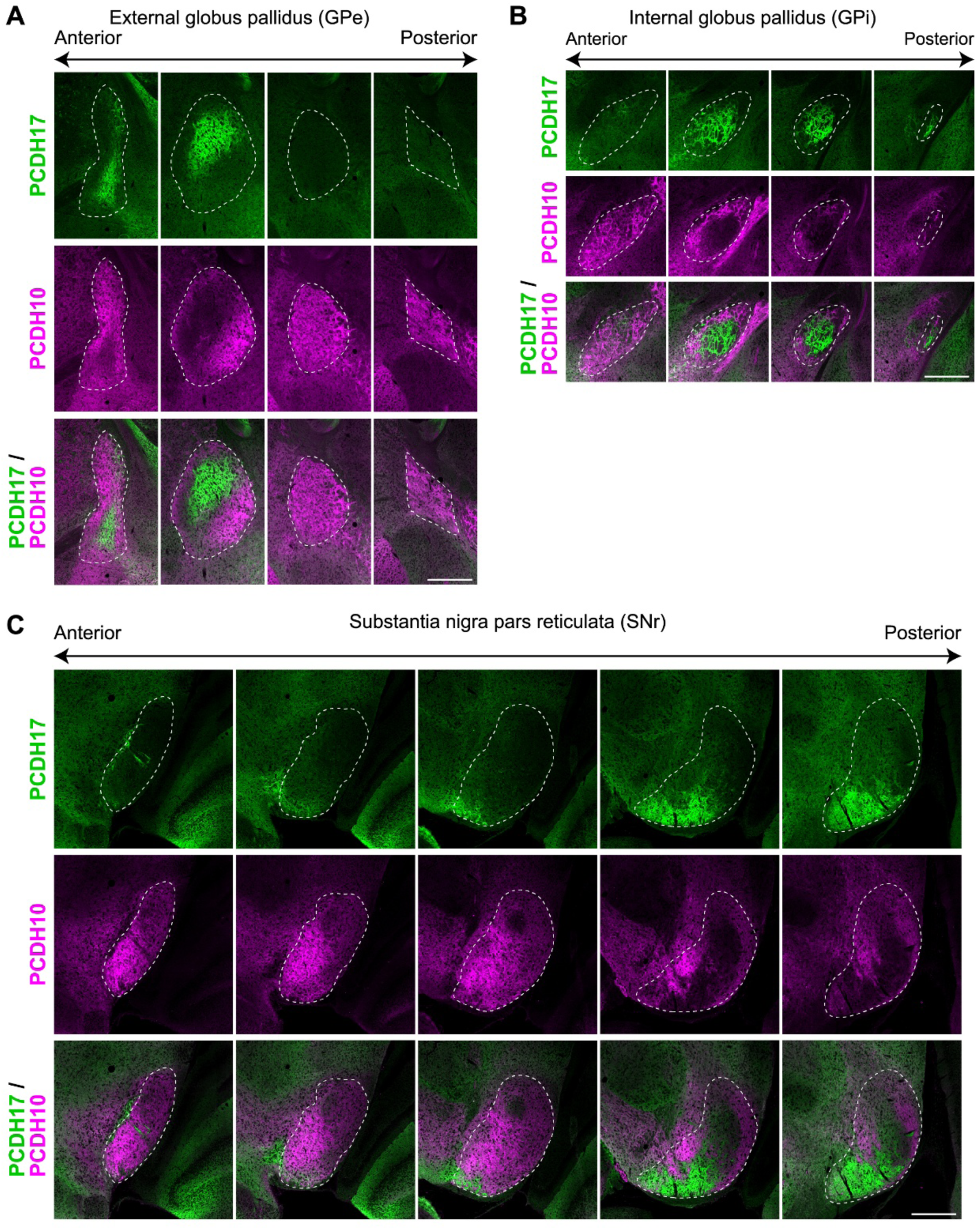
Complementary expression patterns of PCDH17 and PCDH10 proteins in the GPe, GPi, and SNr. Double staining for PCDH17 (green) and PCDH10 (magenta) of coronal BG sections from P11 mice. PCDH17 and PCDH10 proteins show complementary expression patterns in the GPe (**A**), GPi (**B**), and SNr (**C**). The scale bars represent 0.5 mm.

**Fig. S3.**
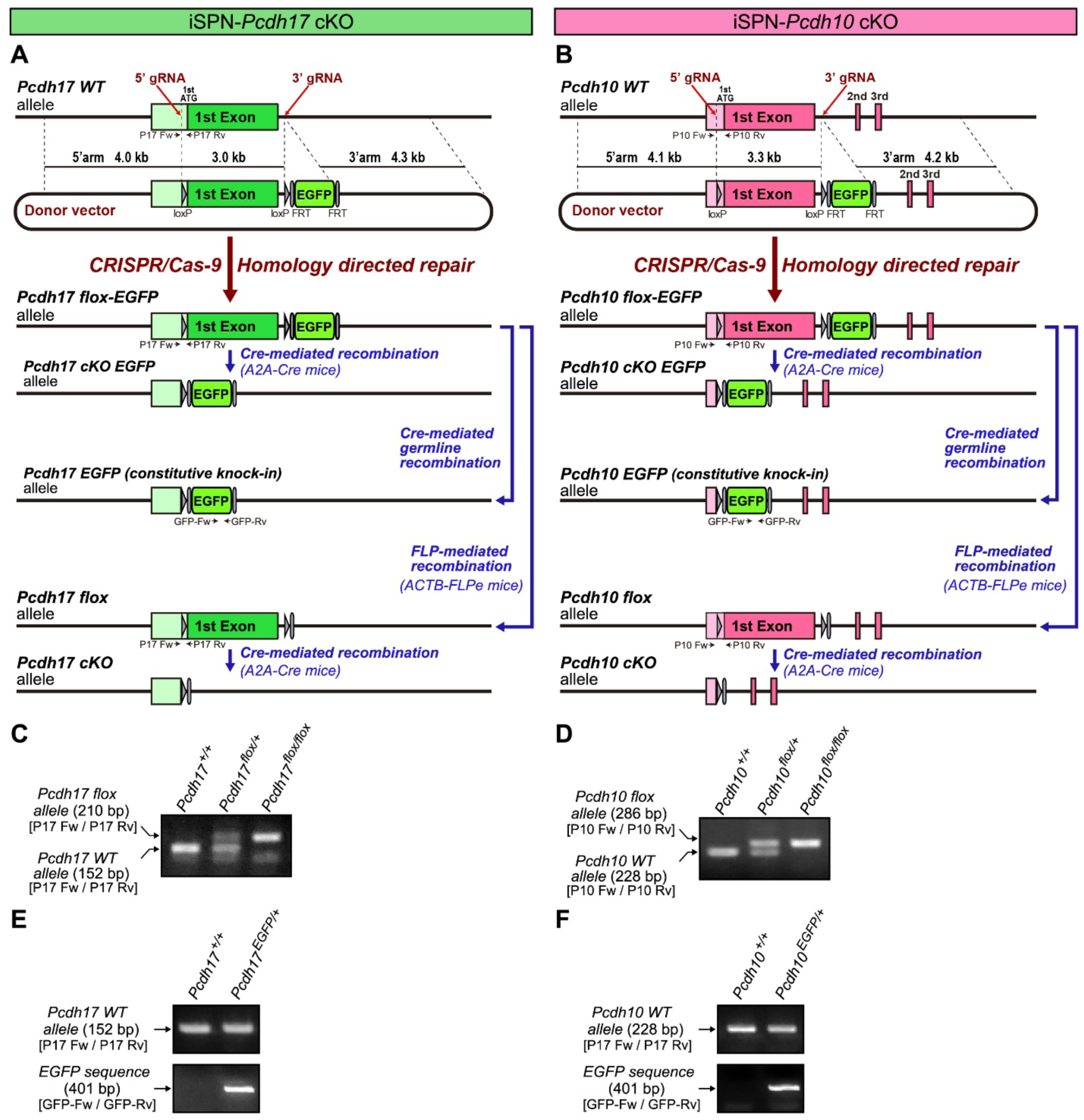
Generation of iSPN-specific cKO mice for *Pcdh17* and *Pcdh10*. (**A** and **B**) Schematic illustrations of the strategy for generating conditional knockout (cKO) mice for *Pcdh17* (**A**) and *Pcdh10* (**B**) using the CRISPR/Cas9 system. For each *Pcdh17* and *Pcdh10* gene: the WT allele, donor vector, flox-EGFP allele, cKO-EGFP allele, EGFP knock-in allele, flox allele, and cKO allele are represented. The donor vector contained the loxP-flanked 1st exon and the FRT-flanked EGFP cassette, along with ∼4 kb of 5’ and 3’ homology arms. Two gRNAs, 5’ and 3’, are designed to remove the 1st exon from the WT allele and replace it with the sequence from the donor vector by homology-directed repair, which generates the *Pcdh*-flox-EGFP allele. The iSPN-specific *Pcdh*-cKO-EGFP allele was produced by mating *Pcdh*-flox-EGFP mice with *A2A-Cre* mice. In *Pcdh*-cKO-EGFP mice, the *Pcdh* promoter drives EGFP expression in Cre-positive iSPNs. *Pcdh*-EGFP-knock-in mice were generated via Cre-mediated germline recombination. The *Pcdh*-flox-EGFP allele was converted to the *Pcdh*-flox allele via the FRT-FLP recombination system by crossing with *ACTB-FLPe* mice. iSPN-specific *Pcdh*-cKO mice are generated by mating *Pcdh*-flox mice with *A2A-Cre* mice. P17 Fw and P17 Rv are the *Pcdh17* flox mice genotyping primers. P10 Fw and P10 Rv are the *Pcdh10* flox mice genotyping primers. GFP Fw and GFP Rv are genotyping primers for the EGFP cassette. (C) Genotyping PCR results for *Pcdh17^+/+^*, *Pcdh17^flox/+^*, and *Pcdh17^flox/flox^* mice. (D) Genotyping PCR results for *Pcdh10^+/+^*, *Pcdh10^flox/+^*, and *Pcdh10^flox/flox^* mice. (E) Genotyping PCR results for *Pcdh17^+/+^* and *Pcdh17^EGFP/+^* mice. (F) Genotyping PCR results for *Pcdh10^+/+^* and *Pcdh10^EGFP/+^* mice.

**Fig. S4.**
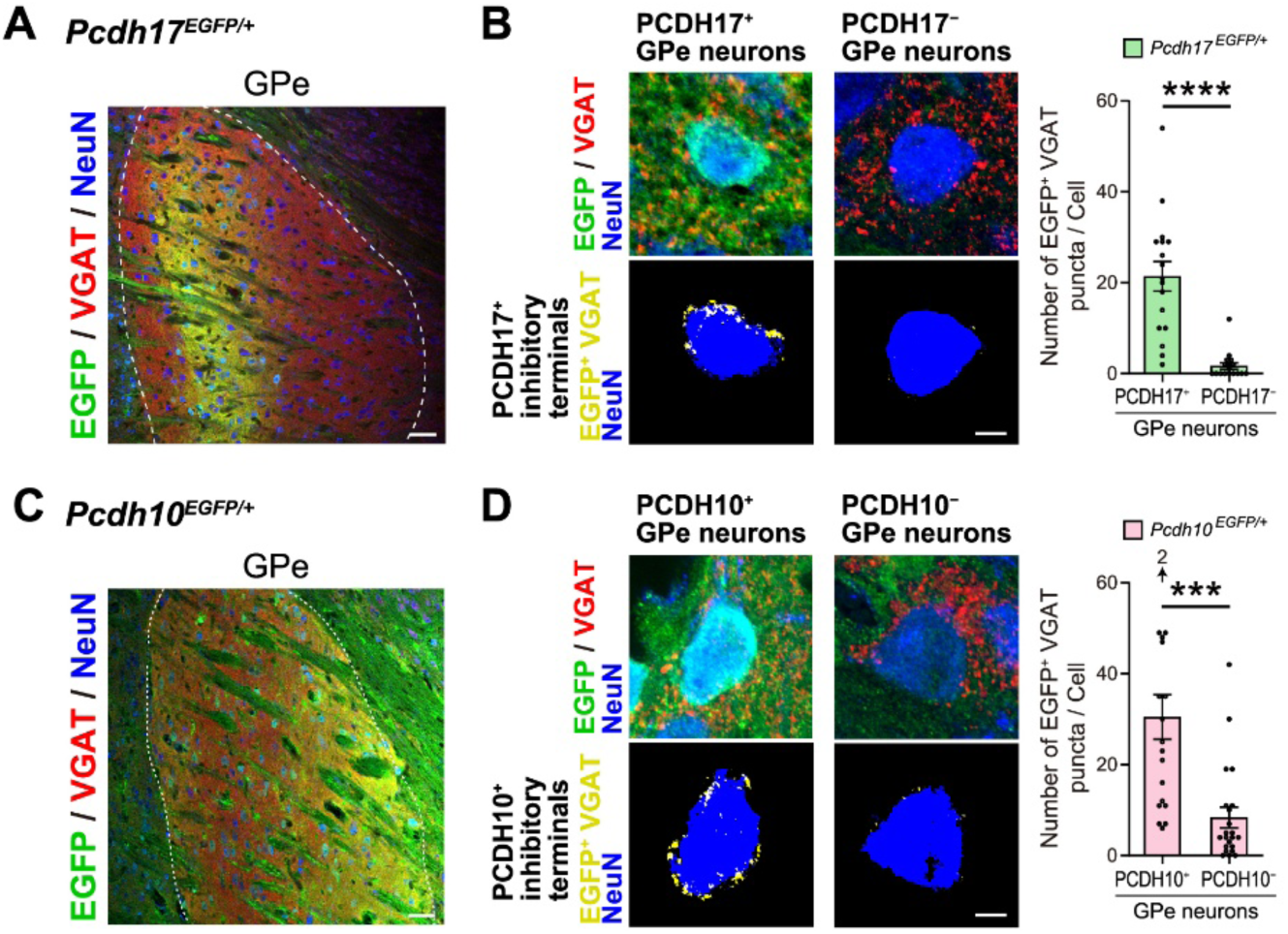
PCDH17- and PCDH10-expressing iSPNs innervate GPe neurons with matching PCDH expression. (**A**) Sagittal sections of the GPe from P11 *Pcdh17^EGFP/+^* mice were immunostained for EGFP (green), VGAT (red), and NeuN (blue). EGFP labels both PCDH17^+^ GPe neurons and PCDH17^+^ inhibitory terminals. (**B**) PCDH17⁺ inhibitory terminals targeting the cell body of PCDH17^+^ GPe neurons (EGFP⁺/NeuN⁺) and PCDH17^–^ GPe neurons (EGFP⁻/NeuN⁺). PCDH17⁺ inhibitory terminals (EGFP⁺/VGAT⁺ yellow puncta) were extracted by overlaying binarized EGFP and VGAT signals. NeuN signals were also binarized to identify cell bodies (blue). Quantification revealed that PCDH17⁺ inhibitory terminals selectively innervate PCDH17^+^ GPe neurons. *n* = 17 cells. (**C**) Sagittal sections of the GPe from P13 *Pcdh10^EGFP/+^* mice were immunostained for EGFP (green), VGAT (red), and NeuN (blue). EGFP labels both PCDH10^+^ GPe neurons and PCDH10^+^ inhibitory terminals. (D) PCDH10⁺ inhibitory terminals targeting the cell bodies of PCDH10^+^ GPe neurons (EGFP⁺/NeuN⁺) and PCDH10^–^ GPe neurons (EGFP⁻/NeuN⁺). PCDH10⁺ inhibitory terminals (EGFP⁺/VGAT⁺ yellow puncta) were extracted by overlaying binarized EGFP and VGAT signals. NeuN signals were also binarized to identify cell bodies (blue). Quantification revealed that PCDH10⁺ inhibitory terminals preferentially innervate PCDH10^+^ GPe neurons. *n* = 19-22 cells. The scale bars represent 50 μm (**a,b**) and 5 μm (**c,d**). Data are mean ± SEM. \*\*\**P* < 0.001, \*\*\*\**P* < 0.0001 by Student’s *t*-test.

**Fig. S5.**
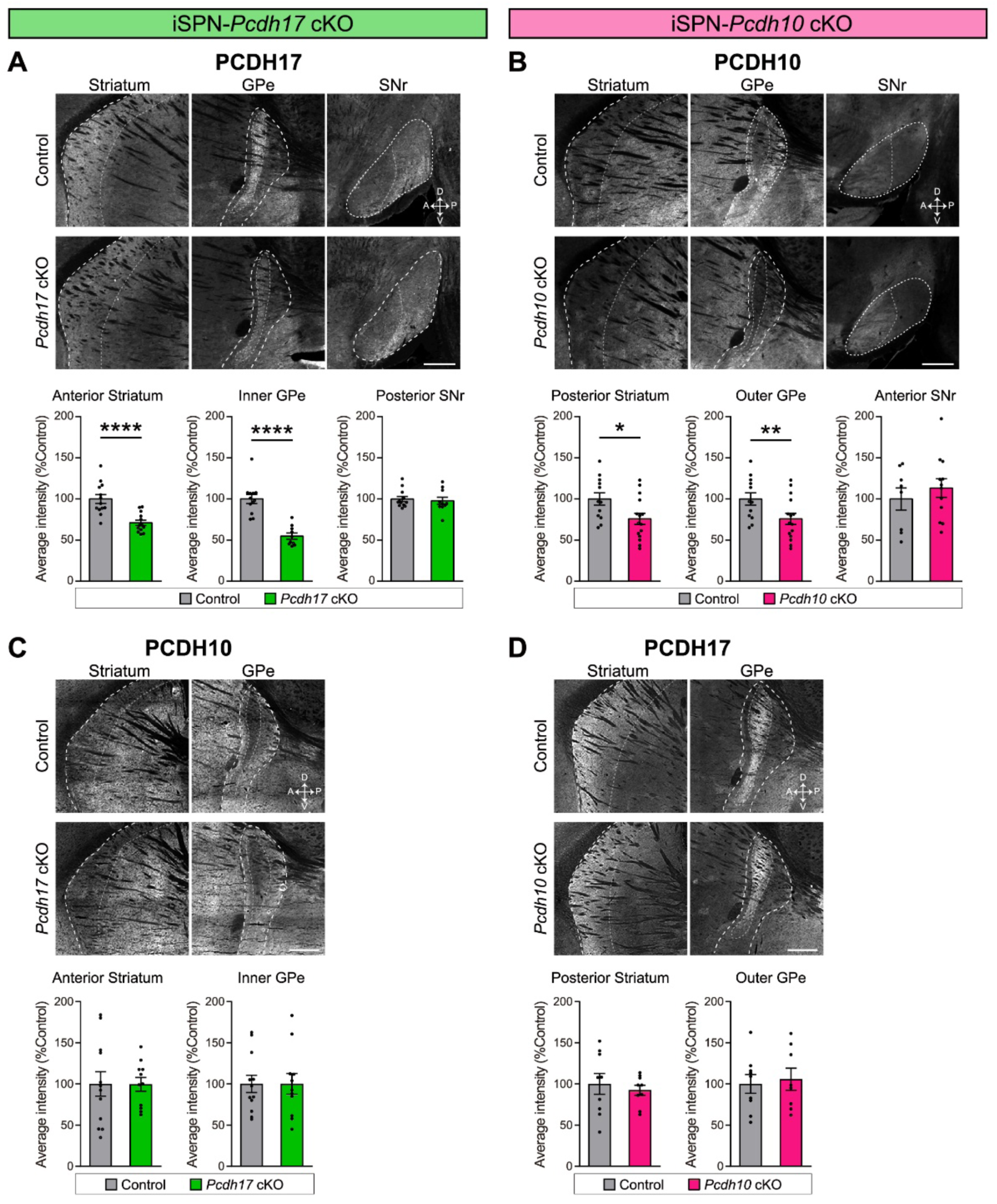
Conditional knockout of PCDH17 or PCDH10 in indirect BG circuits. (**A**) (Top) Immunostaining for PCDH17 of BG sections from adult control (*A2A-Cre;Pcdh17^+/+^*) and *Pcdh17* cKO (*A2A-Cre;Pcdh17^flox/flox^*) mice. (Bottom) Average intensities of PCDH17 staining signals in the anterior striatum, inner GPe, and posterior SNr were quantified. *Pcdh17* cKO mice show reduced PCDH17 staining intensity in the striatum and GPe, but not in the SNr, compared to control mice. n = 10–13 fields from 3 mice per genotype. (**B**) (Top) Immunostaining for PCDH10 of BG sections from adult control (*A2A-Cre;Pcdh10^+/+^*) and *Pcdh10* cKO (*A2A-Cre;Pcdh10^flox/flox^*) mice. (Bottom) Average intensities of PCDH10 staining signals in the posterior striatum, outer GPe, and anterior SNr were quantified. *Pcdh10* cKO mice show reduced PCDH10 staining intensity in the striatum and GPe, but not in the SNr, compared to control mice. n = 8–15 fields from 4–5 mice per genotype. (**C**) (Top) Immunostaining for PCDH10 of BG sections from 3-week-old control and *Pcdh17* cKO mice. (Bottom) Average intensities of PCDH10 staining signals in the anterior striatum and inner GPe were quantified. *Pcdh17* cKO mice show similar levels of PCDH10 staining intensity in the striatum and GPe compared to control mice. n = 11–12 fields from 6 mice per genotype. (**D**) (Top) Immunostaining for PCDH17 of BG sections from 3-week-old control and *Pcdh10* cKO mice. (Bottom) Average intensities of PCDH17 staining signals in the posterior striatum and outer GPe were quantified. *Pcdh10* cKO mice show similar levels of PCDH17 staining intensity in the striatum and GPe compared to control mice. n = 8–9 fields from 5 mice per genotype. The scale bars represent 0.5 mm. Data are mean ± SEM. \**P* < 0.05, \*\**P* < 0.01, \*\*\*\**P* < 0.0001 by Student’s *t*-test. GPe, external globus pallidus; GPi, internal globus pallidus; SNr, substantia nigra pars reticulata.

**Fig. S6.**
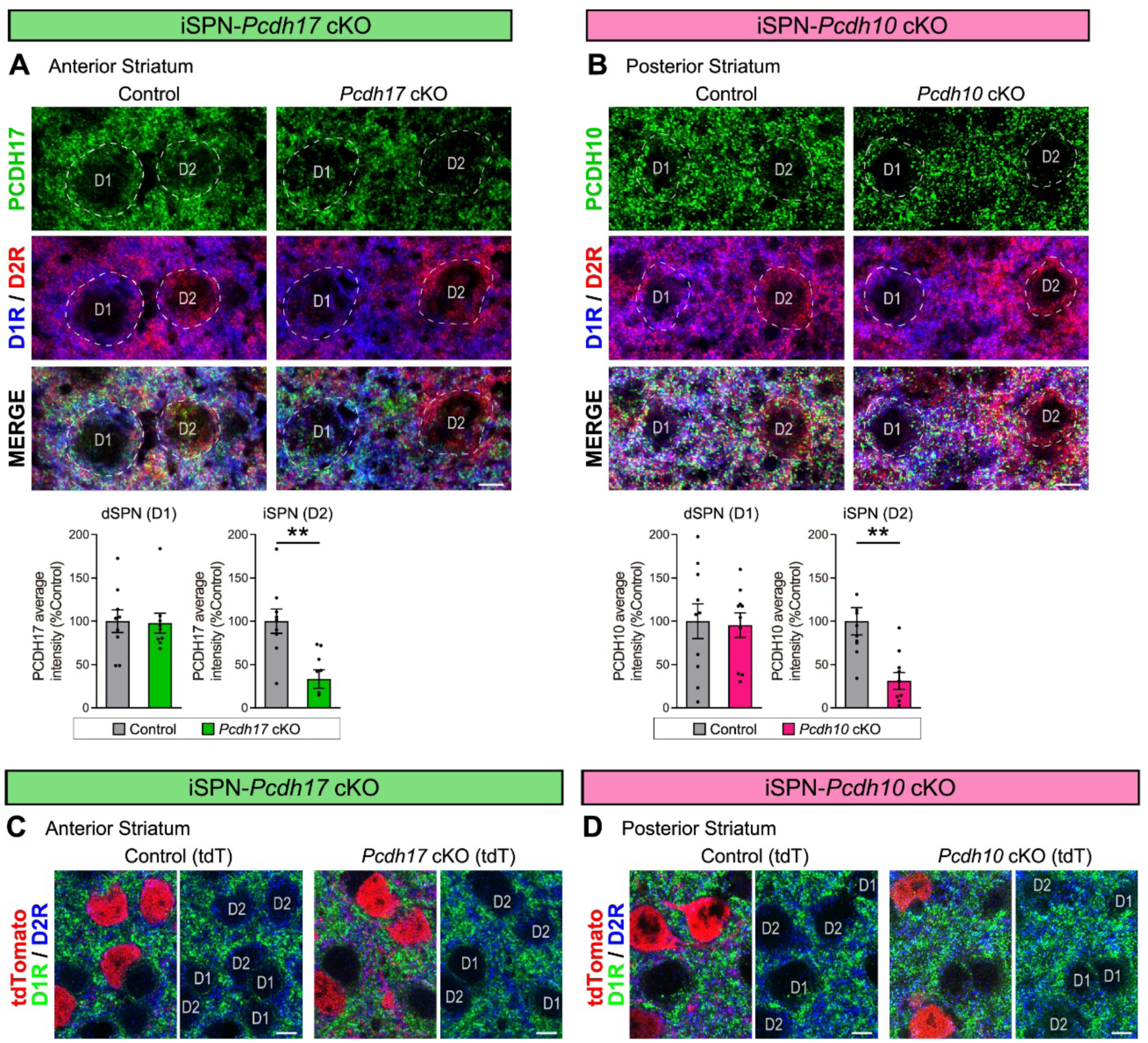
iSPN-specific knockout of PCDH17 or PCDH10 in the striatum. (**A**) Triple staining for PCDH17 (green), D1R (blue), and D2R (red) in the anterior striatum of 3-week-old control and *Pcdh17* cKO mice. Average intensities of PCDH17 staining signals on the cell bodies of dSPNs (D1R-positive) and iSPNs (D2R-positive) were quantified. *Pcdh17* cKO mice show significantly reduced PCDH17 staining signals in iSPNs but not in dSPNs compared to control mice. *n* = 9 cells for iSPNs and dSPNs from 5 mice per genotype. (**B**) Triple staining for PCDH10 (green), D1R (blue), and D2R (red) in the posterior striatum of 3-week-old control and *Pcdh10* cKO mice. Average intensities of PCDH10 staining signals on the cell bodies of dSPNs and iSPNs were quantified. *Pcdh10* cKO mice show significantly reduced PCDH10 staining signals in iSPNs but not in dSPNs compared to control mice. *n* = 10 cells for iSPNs and dSPNs from 5 mice per genotype. (**C** and **D**) iSPN cells were labeled with tdTomato by crossing *A2A-Cre* mice with *Rosa^STOP-tdT^* mice. Staining for D1R (green) and D2R (blue) in the anterior striatum of 3-week-old control (tdT) and *Pcdh17* cKO (tdT) mice (**C**) and in the posterior striatum of 3-week-old control (tdT) and *Pcdh10* cKO (tdT) mice (**D**). tdTomato-positive iSPN cells are D2R-positive across all genotypes. The scale bars represent 5 μm. Data are mean ± SEM. \*\**P* < 0.01 by Student’s *t*-test.

**Fig. S7.**
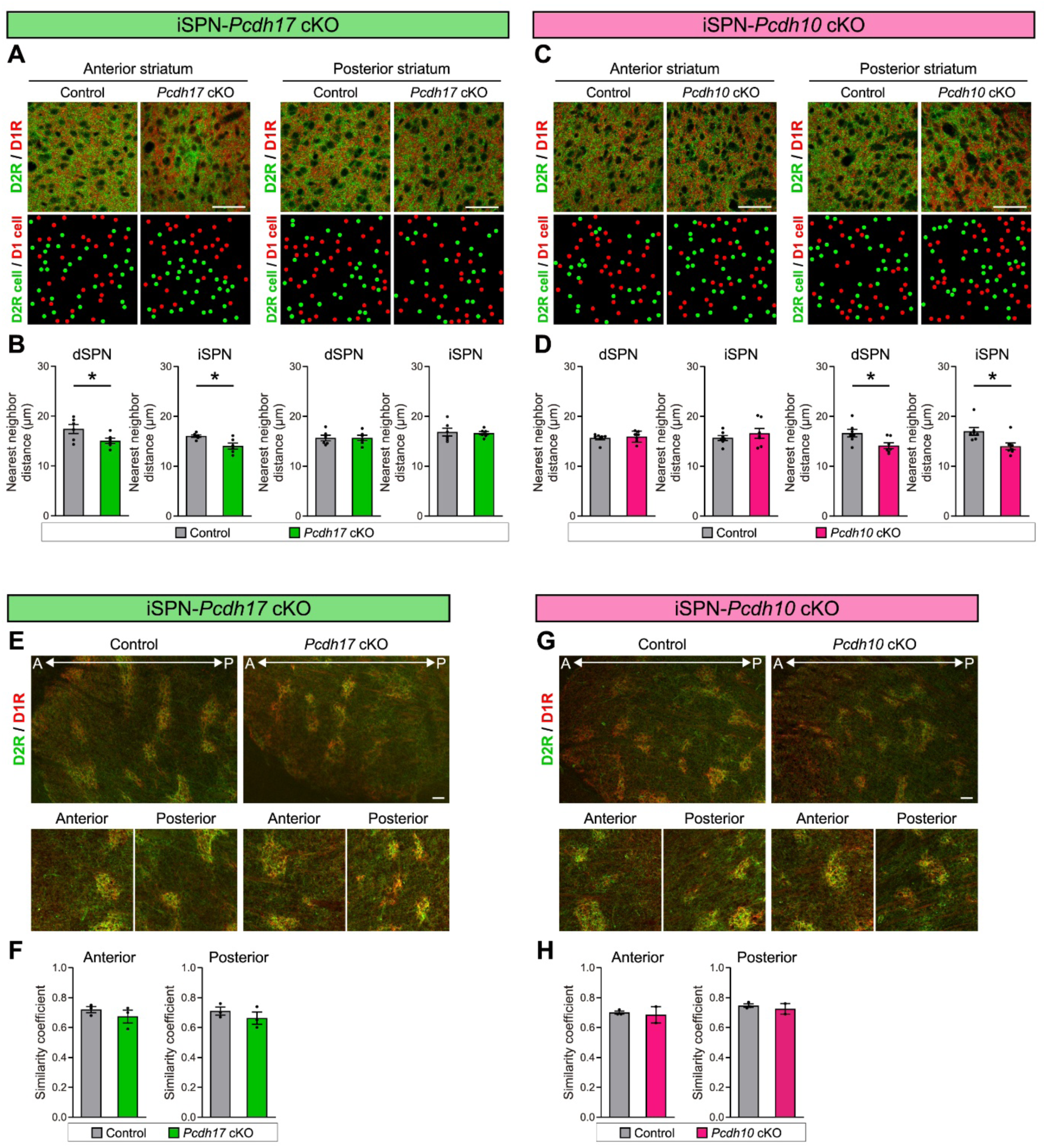
Abnormal cell body spacing of SPNs in the striatum of 3-week-old iSPN-*Pcdh* cKO mice and normal D1R and D2R distribution in the striatum of iSPN-*Pcdh* cKO mice at postnatal day 0. (**A** to **D**) Region-specific abnormal cell body spacing of dSPNs (D1R-positive cells) and iSPNs (D2R-positive cells) in iSPN-*Pcdh* cKO mice. (**A**) (Top) Double staining for D1R (red; labels dSPNs) and D2R (green; labels iSPNs) of anterior and posterior striatal sections from 3-week-old *Pcdh17* cKO and control mice. (Bottom) The positions of dSPN and iSPN cell bodies were extracted and marked with red and green circles, respectively. (**B**) Quantification of the nearest neighbor distance of dSPN and iSPN cell bodies in the anterior and posterior striatum. *n* = 6 (total) fields from 3 mice per genotype. (**C**) (Top) Double staining for D1R (red; labels dSPNs) and D2R (green; labels iSPNs) of anterior and posterior striatal sections from 3-week-old *Pcdh10* cKO and control mice. (Bottom) The positions of dSPN and iSPN cell bodies were extracted and marked with red and green circles, respectively. (**D**) Quantification of the nearest neighbor distance of dSPN and iSPN cell bodies in the anterior and posterior striatum. *n* = 7 (total) fields from 4 mice per genotype. (**E** to **H)** Normal D1R and D2R distribution in postnatal day 0 (P0) iSPN-*Pcdh* cKO mice. (**E**) Double staining for D2R (green; iSPNs) and D1R (red; dSPNs) of anterior and posterior striatal sections from P0 *Pcdh17* cKO and control mice. (**F**) Quantification of the similarity coefficient in the anterior and posterior striatum. *n* = 3 fields from 3 mice per genotype. (**G**) Double staining for D1R and D2R of anterior and posterior striatal sections from P0 *Pcdh10* cKO and control mice. (**H**) Quantification of the similarity coefficient in the anterior and posterior striatum. *n* = 3 fields from 3 mice per genotype. The scale bars represent 50 μm. Data are mean ± SEM. \**P* < 0.05 by Student’s *t*-test.

**Fig. S8:**
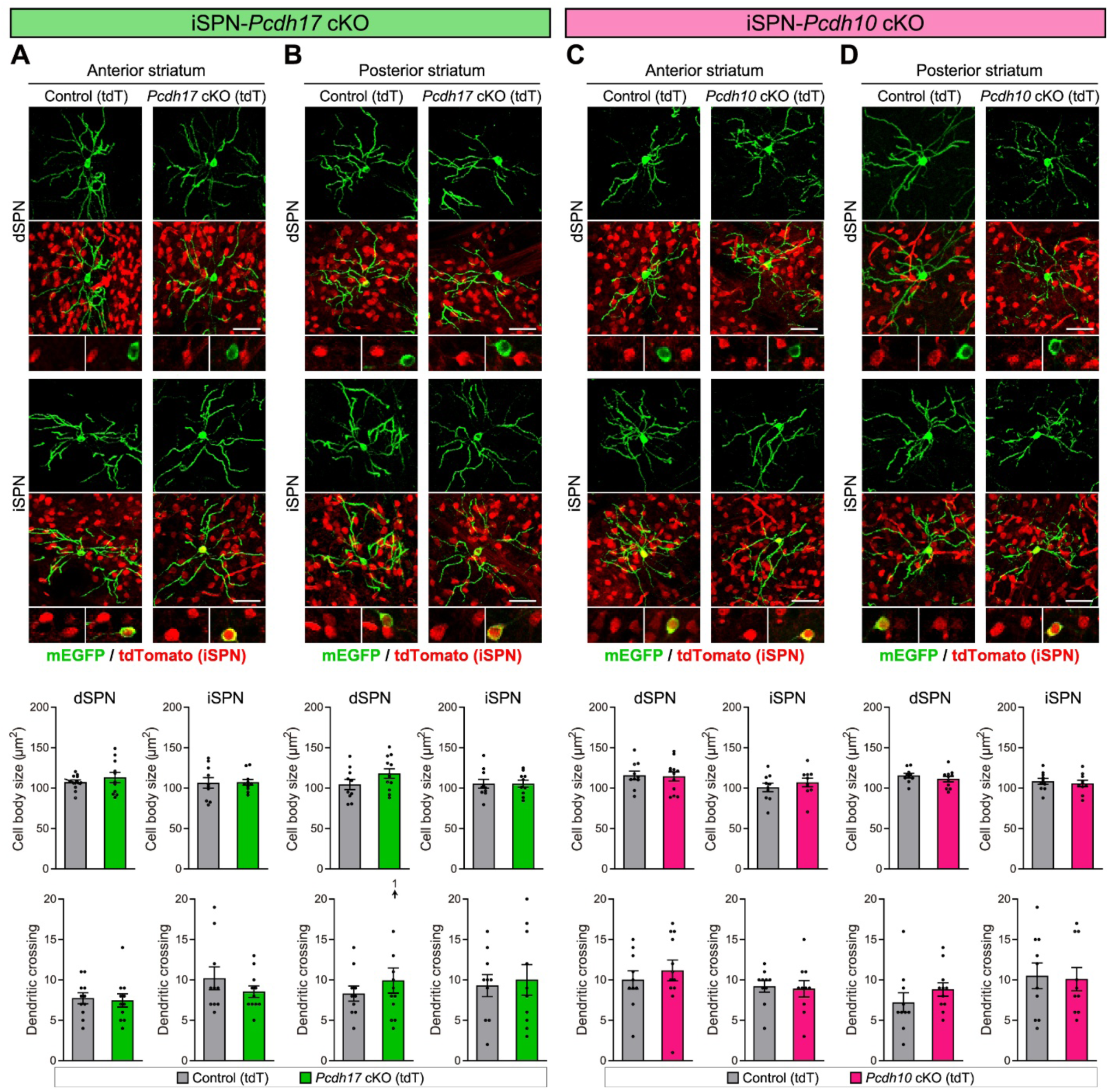
The morphology of dSPNs and iSPNs is normal in iSPN-*Pcdh* cKO mice. (**A** to **D**) The morphology of dSPNs and iSPNs, assessed by labeling with membrane-EGFP (mEGFP), in the anterior and posterior striatum of adult *Pcdh* cKO (tdT) and control mice (tdT). tdTomato^+^ cells are iSPNs. Single-plane images of the cell bodies are magnified in the bottom panels to demonstrate mEGFP^+^/tdTomato^-^ cells (dSPNs) and mEGFP^+^/tdTomato^+^ cells (iSPNs). (**A** and **B**) The cell body size and number of dendritic crossings of dSPNs and iSPNs in both the anterior (**A**) and posterior (**B**) striatum are similar between control (tdT) and *Pcdh17* cKO (tdT) mice. *n* = 10-12 cells for iSPNs and dSPNs in the anterior and posterior striatum from 3 mice per genotype. (**C** and **D**) The cell body size and number of dendritic crossings of dSPNs and iSPNs in both the anterior (**C**) and posterior (**D**) striatum are similar between control (tdT) and *Pcdh10* cKO (tdT) mice. *n* = 10-12 cells for iSPNs and dSPNs in the anterior and posterior striatum from 3 mice per genotype. The scale bars represent 50 μm. Data are mean ± SEM. Student’s *t*-test.

**Fig. S9:**
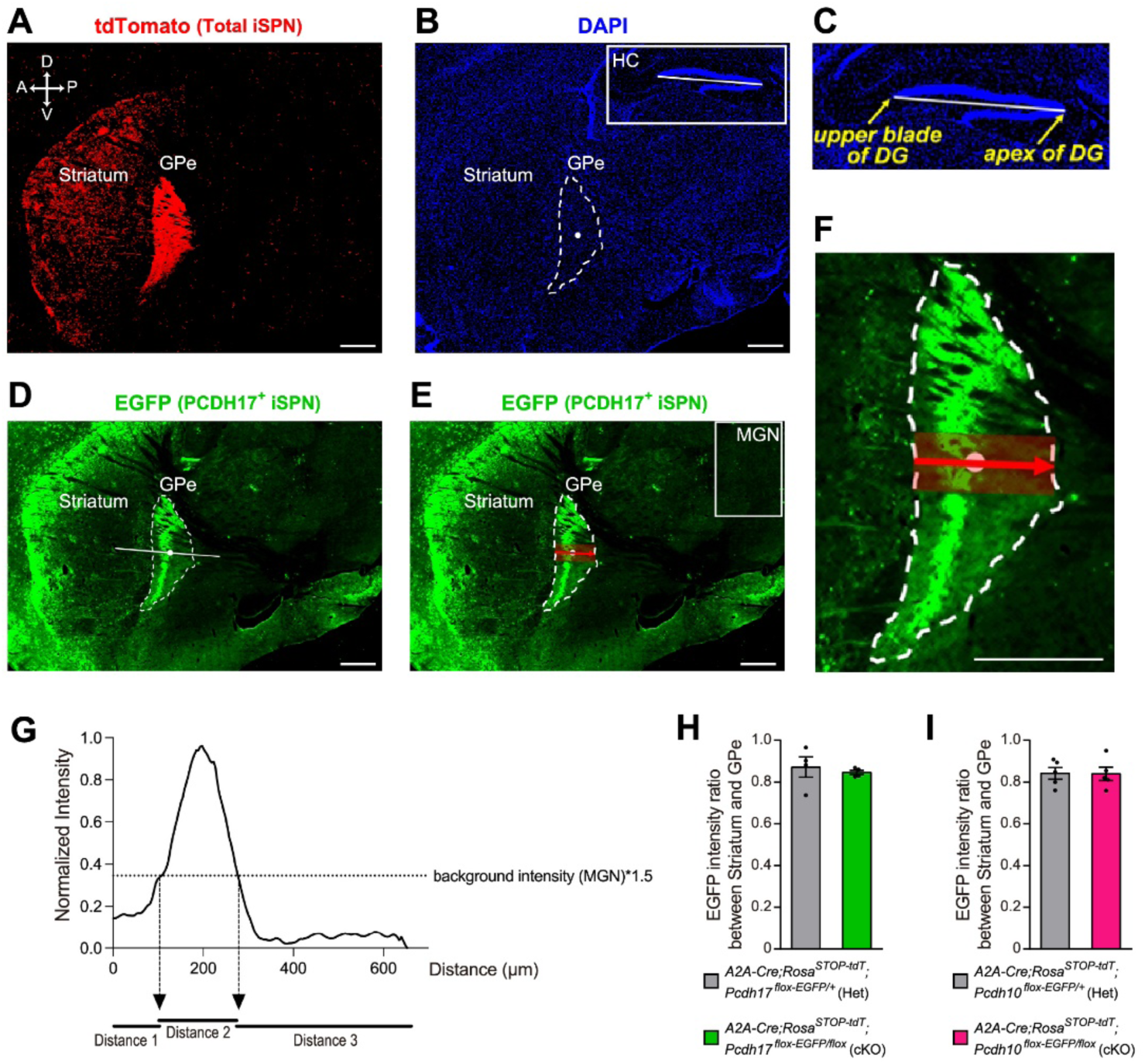
Method for axonal projection analysis using sagittal brain sections from *A2A-Cre;Rosa^STOP-tdT^;Pcdh^flox-EGFP^*mice. (**A** to **G**) Evaluation of iSPN axon targeting in the GPe. (**a**) The tdTomato signal labels all the iSPNs within the striatum and their axonal projections to the GPe in red. The outline of the tdTomato signal in the GPe (dashed, white line) was used to determine the geometrical center of the GPe (white dot within dashed outline). (**B** and **C**) DAPI staining was used to construct a line in the hippocampus (HC, white box in **B**). The line connects the apex of the dentate gyrus (DG) and the anterior end of the upper blade of the DG (**C**). A parallel to this line was then drawn through the geometrical center of the GPe (**B**). (**D**) Immunostaining for EGFP (green) visualizes the area of iSPN axonal projection within the GPe as a result of EGFP expression under the *Pcdh* promoter in iSPNs. (**E** and **F**) The line through the geometrical center was used to perform a 50-pixel wide line scan crossing the GPe from the anterior to the posterior side (the red arrow indicates the orientation of the line scan; width of the line scan is represented by a red box). (**G**) The resulting line scan (the highest intensity in the scan was set as 1 and the remaining values were normalized accordingly) was evaluated. The background intensity was calculated from the mean intensity in the medial geniculate nucleus (MGN, white box in **E**) and introduced in the line scan (dotted line). This resulted in two crossing points with the line scan, which marked the borders of axonal innervation. We then calculated the distances between the beginning of the line scan and the first border of axonal innervation (Distance 1), the first and second borders of axonal innervation (Distance 2), and the second border of axonal innervation and the end of the line scan (Distance 3). (**H** and **I**) Evaluation of iSPN axon density in the GPe. EGFP intensity ratios between the striatum and GPe were calculated. (**H**) Intensity ratio between the anterior striatum and the inner GPe are similar between *A2A-Cre;Rosa^STOP-tdT^;Pcdh17^flox-EGFP/flox^*mice (cKO) and *A2A-Cre;Rosa^STOP-tdT^;Pcdh17^flox-EGFP/+^*mice (Het). (**I**) Intensity ratio between the posterior striatum and the posterior GPe are similar between *A2A-Cre;Rosa^STOP-tdT^;Pcdh10^flox-EGFP/flox^* mice (cKO) and *A2A-Cre;Rosa^STOP-tdT^;Pcdh10^flox-EGFP/+^*mice (Het). The scale bars represent 0.5 mm. Data are mean ± SEM. Student’s *t*-test.

**Fig. S10:**
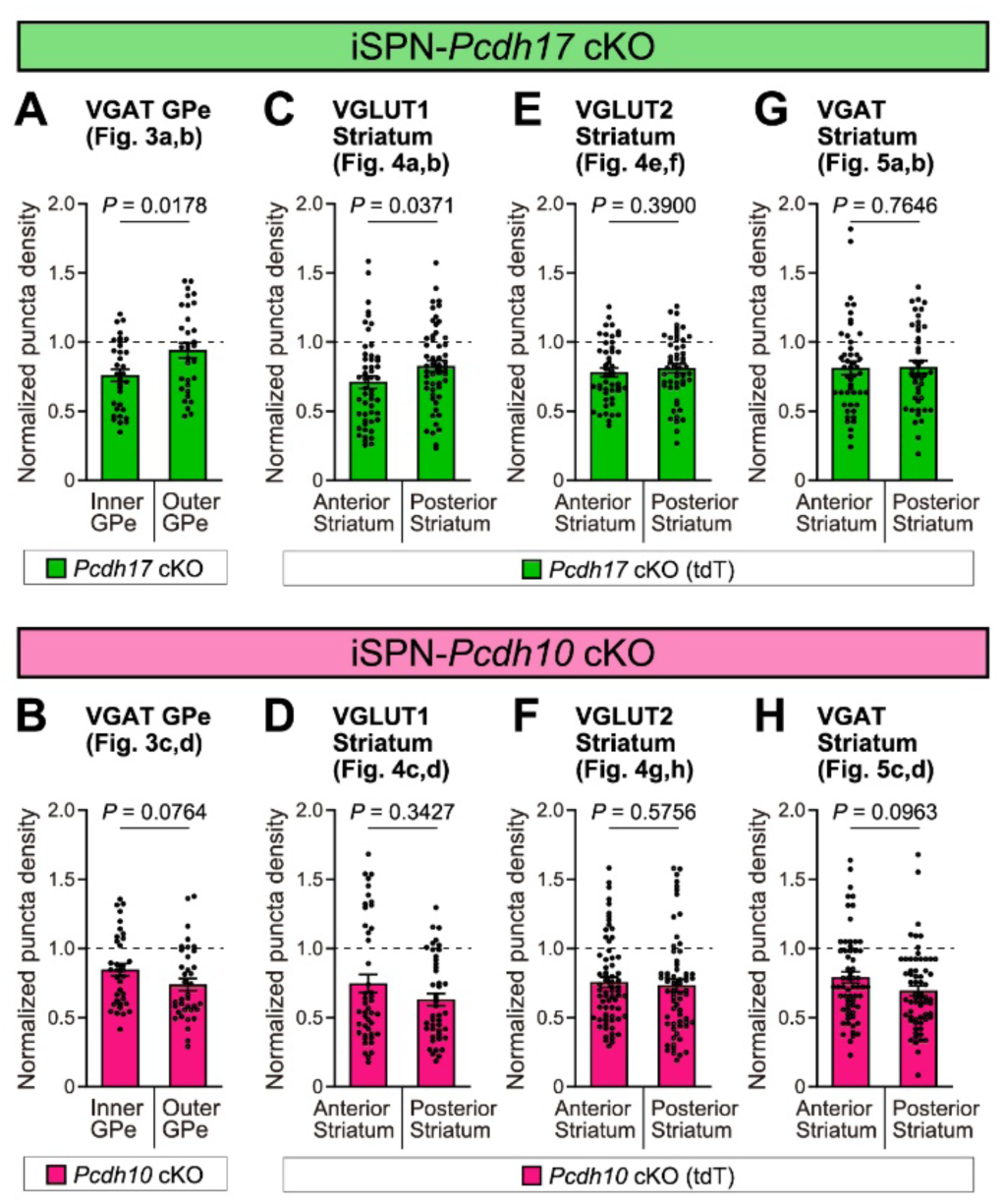
Comparison of normalized synaptic puncta density between the inner and outer GPe, as well as between the anterior and posterior striatum, in iSPN-*Pcdh* cKO mice. (**A** and **B**) Comparison of normalized VGAT puncta density between the inner and outer GPe in *Pcdh17* cKO and *Pcdh10* cKO mice from Fig. 3a-d. (**C** and **D**) Comparison of normalized VGLUT1 puncta density between the anterior and posterior striatum in *Pcdh10* cKO(tdT) and *Pcdh17* cKO (tdT) mice from Fig. 4, A to D. (**E** and **F**) Comparison of normalized VGLUT2 puncta density between the anterior and posterior striatum in *Pcdh10* cKO (tdT) and *Pcdh17* cKO (tdT) mice from Fig. 4, E to H. (**G** and **H**) Comparison of normalized VGAT puncta density between the anterior and posterior striatum in *Pcdh10* cKO (tdT) and *Pcdh17* cKO (tdT) mice from Fig. 5, A to D. Data are mean ± SEM. Mann-Whitney U tests were performed between different regions in iSPN-*Pcdh* cKO mice, and the *P*-values are indicated.

**Fig. S11:**
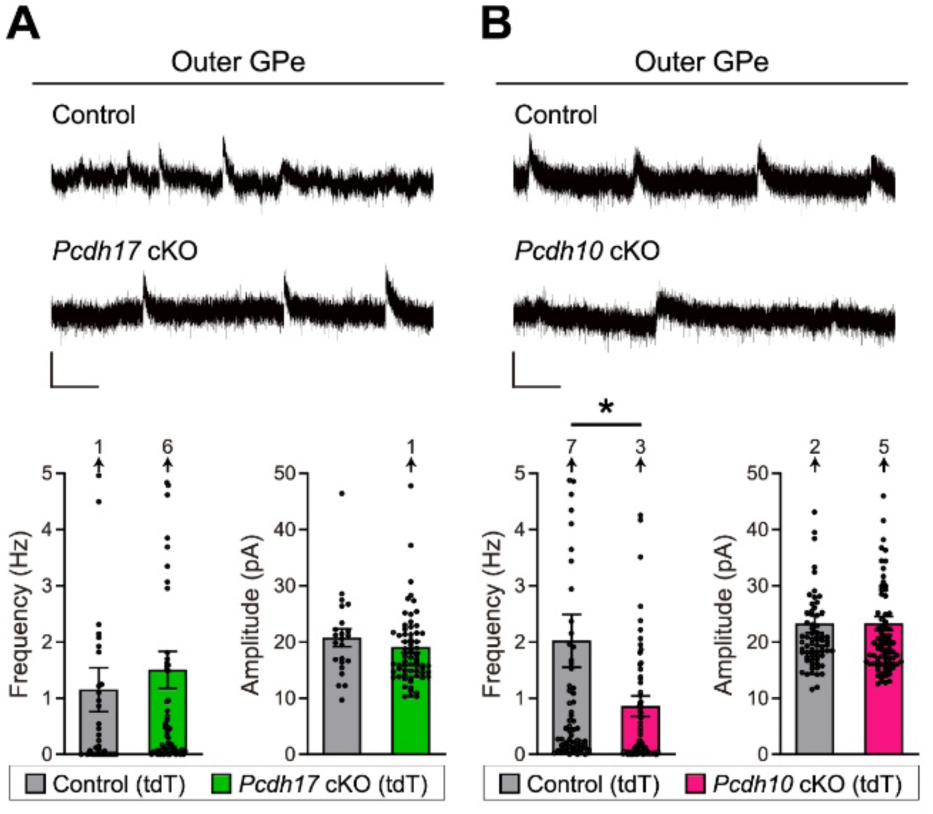
*Pcdh10* cKO, but not *Pcdh17* cKO mice, show reduced sIPSC frequency recorded from outer GPe neurons. (**A** and **B**) Spontaneous IPSCs (sIPSCs) recorded from outer GPe neurons. The top panels show example traces. The graphs show the frequency and amplitude of sIPSCs in *Pcdh* cKO (tdT) and control (tdT) mice. (**A**) Both the frequency and the amplitude of sIPSCs are similar between *Pcdh17* cKO (tdT) and control (tdT) mice. (**B**) The frequency, but not the amplitude, of sIPSCs is decreased in *Pcdh10* cKO (tdT) mice compared to control (tdT) mice. Frequency: *n* = 34 neurons from 4 *Pcdh17* control (tdT) mice; 72 neurons from 10 *Pcdh17* cKO (tdT) mice; 71 neurons from 8 *Pcdh10* control (tdT) mice; and 99 neurons from 12 *Pcdh10* cKO (tdT) mice. Amplitude: *n* = 23 neurons from 4 *Pcdh17* control (tdT) mice; 66 neurons from 10 *Pcdh17* cKO (tdT) mice; 68 neurons from 8 *Pcdh10* control (tdT) mice; and 85 neurons from 12 *Pcdh10* cKO (tdT) mice. The scale bars represent 20 pA and 100 ms. Data are mean ± SEM. \**P* = 0.0128, Student’s *t*-test.

**Fig. S12:**
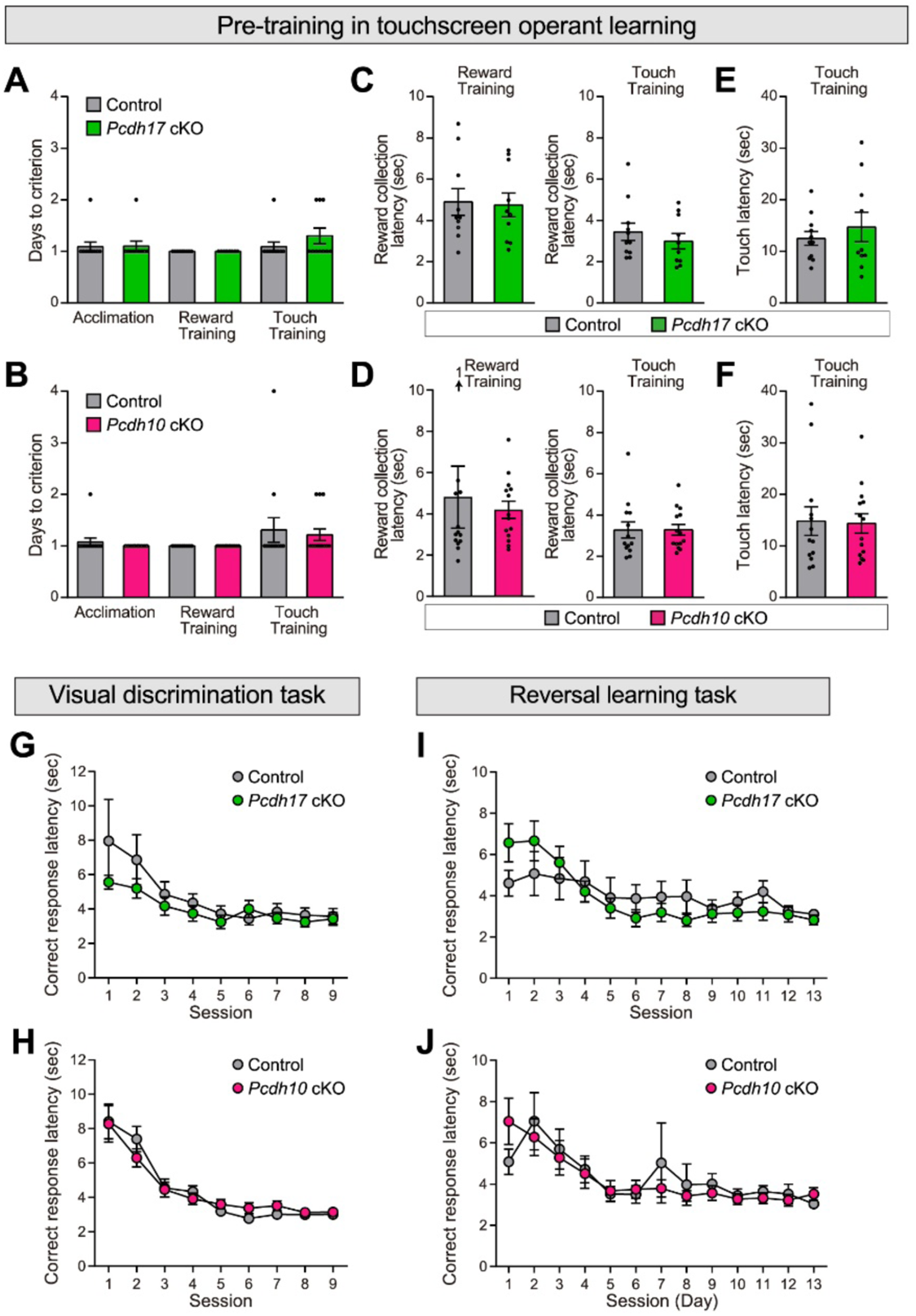
Parameters during pre-training and correct response latencies during visual discrimination and reversal learning tasks are normal in *Pcdh* cKO mice. (**A** to **F**) Pre-training. (**A** and **B**) Days to criterion in acclimation, reward acquisition training, and stimulus touch training are similar between *Pcdh17* cKO mice and control mice (**A**) and between *Pcdh10* cKO mice and control mice (**B**). (**C** and **D**) Reward latencies during reward training (left) and touch training (right) are similar between *Pcdh* cKO mice and control mice. (**E** and **F**) Touch latencies during touch training are similar between *Pcdh* cKO mice and control mice. n = 11 mice for *Pcdh17* control; 10 for *Pcdh17* cKO; 13 for *Pcdh10* control; and 14 for *Pcdh10* cKO. (**G** and **H**) Correct response latencies during the visual discrimination task are similar between *Pcdh17* cKO mice and *Pcdh17* control mice (**G**) and between *Pcdh10* cKO mice and *Pcdh10* control mice (**H**). *n* = 10 mice for *Pcdh17 control*; 10 for *Pcdh17 cKO;* 13 for *Pcdh10 control*; and 14 for *Pcdh10 cKO*. (**I** and **J**) Correct response latencies during the reversal learning task are similar between *Pcdh17* cKO mice and control mice (**I**) and between *Pcdh10* cKO mice and control mice (**J**). *n* = 10 mice for *Pcdh17 control*; 10 for *Pcdh17 cKO;* 13 for *Pcdh10 control*; and 14 for *Pcdh10 cKO*. Data are mean ± SEM. Student’s *t*-test.

**Fig. S13:**
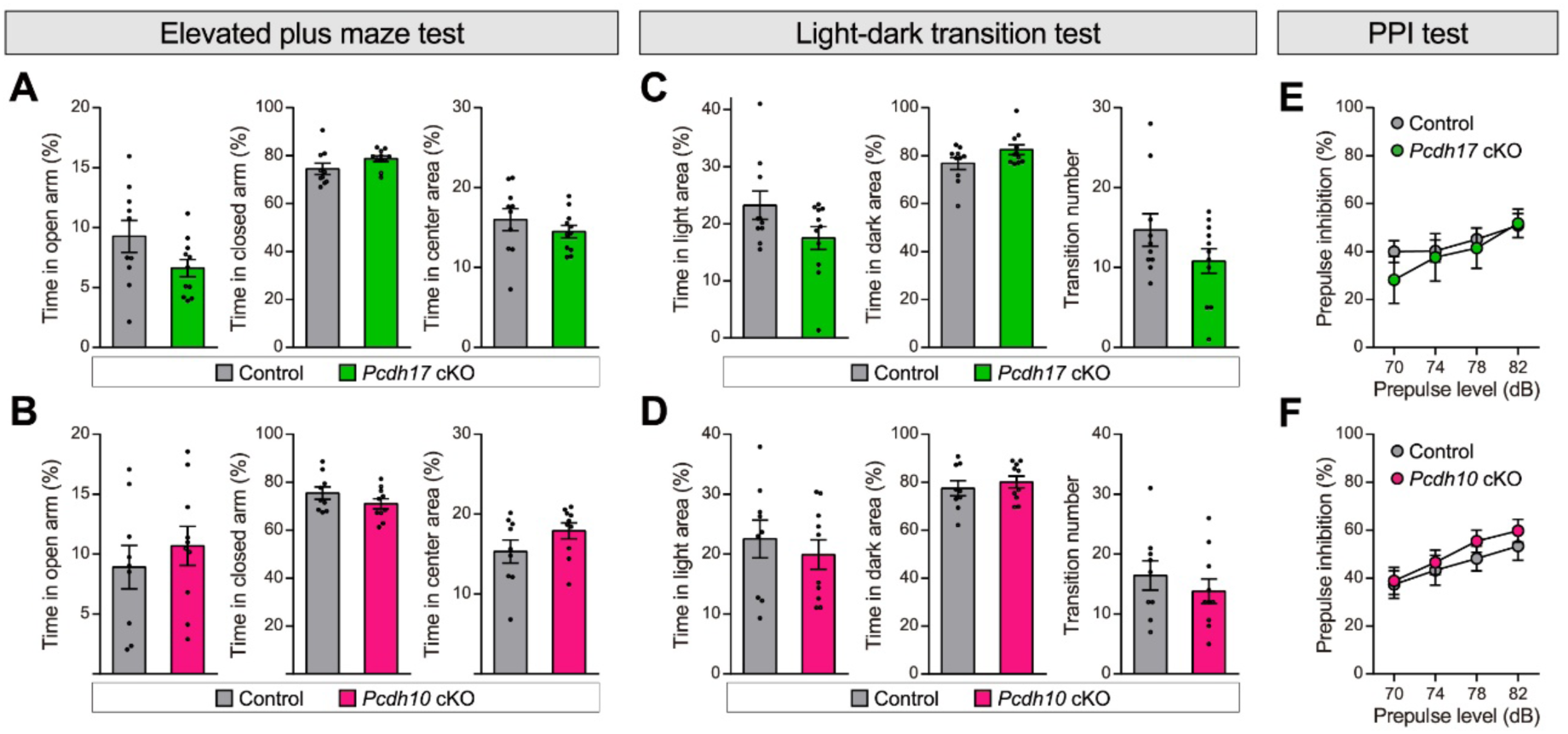
Normal anxiety levels and sensorimotor gating in *Pcdh* cKO mice. (**A** and **B**) Elevated plus maze test. Times spent in the open arm, closed arm, and center area were recorded. There is no significant change in the times spent in each area between *Pcdh* cKO and control mice. *n* = 10 mice for *Pcdh17 control*; 11 for *Pcdh17 cKO;* 9 for *Pcdh10 control*; and 10 for *Pcdh10 cKO*. (**C** and **D**) Light-dark transition test. Times spent in the light and dark areas as well as transition numbers were quantified. There is no significant difference in these measures between *Pcdh cKO* mice and c*ontrol* mice. *n* = 10 mice for *Pcdh17 control*; 11 for *Pcdh17 cKO;* 9 for *Pcdh10 control*; and 10 for *Pcdh10 cKO*. (**E** and **F**) PPI test. PPI (%) at different prepulse levels is similar between *Pcdh17 control* and *Pcdh17 cKO* mice (**E**) as well as between *Pcdh10 control* and *Pcdh10 cKO* mice (**F**). *n* = 11 mice for *Pcdh17 control*; 9 for *Pcdh17 cKO;* 11 for *Pcdh10 control*; and 11 for *Pcdh10 cKO*. Data are mean ± SEM. \**P* < 0.05 by Student’s *t* test.

## Notes

### Competing Interest Statement

The authors have declared no competing interest.

